# A Survey on Methods for Predicting Polyadenylation Sites from DNA Sequences, Bulk RNA-seq, and Single-cell RNA-seq

**DOI:** 10.1101/2022.07.17.500329

**Authors:** Wenbin Ye, Qiwei Lian, Congting Ye, Xiaohui Wu

## Abstract

Alternative polyadenylation (APA) plays important roles in modulating mRNA stability, translation, and subcellular localization, and contributes extensively to shaping eukaryotic transcriptome complexity and proteome diversity. Identification of poly(A) sites (pAs) on a genome-wide scale is a critical step toward understanding the underlying mechanism of APA-mediated gene regulation. A number of established computational tools have been proposed to predict pAs from diverse genomic data. Here we provided an exhaustive overview of computational approaches for predicting pAs from DNA sequences, bulk RNA-seq data, and single-cell RNA-seq (scRNA-seq) data. Particularly, we examined several representative tools using RNA-seq and scRNA-seq data from peripheral blood mononuclear cells and put forward operable suggestions on how to assess the reliability of pAs predicted by different tools. We also proposed practical guidelines on choosing appropriate methods applicable to diverse scenarios. Moreover, we discussed in depth the challenges in improving the performance of pA prediction and benchmarking different methods. Additionally, we highlighted outstanding challenges and opportunities using new machine learning and integrative multi-omics techniques and provided our perspective on how computational methodologies might evolve in the future for non-3’ UTR, tissue-specific, cross-species, and single-cell pA prediction.

## Introduction

Precursor mRNA (pre-mRNA) polyadenylation is an essential two-step event in the post-transcriptional regulation of gene expression, which involves the cleavage of the pre-mRNA at the poly(A) site (pA) followed by the addition of an untemplated stretch of adenosines [1, 2]. The selective use of pAs of a single gene, termed alternative polyadenylation (APA), can generate a diversity of isoforms with different 3’ ends and/or encode distinct proteins [3, 4]. APA plays important roles in modulating mRNA stability, translation, and subcellular localization, which contributes extensively to shaping eukaryotic transcriptome complexity and proteome diversity. APA is a widespread regulatory mechanism in eukaryotes, which has been observed in more than 70% of mammalian and plant genes [5–11]. APA is highly tissue specific and dynamically modulated in various conditions, cell types, and/or states [2, 12]. Specific APA programs have been implicated in diverse biological processes and diseases, such as cell activation, proliferation, neurodegenerative disorders, and cancer [3, 4, 13–20]. Given the functional significance of APA, identification and/or quantification of pAs on a genome-wide scale is crucial and may be the first step in understanding the underlying mechanism of APA-mediated gene regulation.

Early studies, dating back to the 1990s, predict pAs using conventional machine learning (ML) models like support vector machine (SVM) [21–25], which distinguish whether a nucleotide sequence contains a pA using a variety of hand-crafted features (**Figure 1**A). In recent years, deep learning (DL) models [26–29] have been shown to provide better performance than traditional ML methods, owing to their great ability for direct and automatic feature extraction and high scalability with large amount of genomic data (Figure 1B). With the advance of next generation sequencing (NGS) technologies, experimental protocols have been designed to capture 3’ ends of mRNAs for direct profiling of genome-wide pAs (Figure 1C), such as DRS [10, 30], 3P-Seq [7, 31], 3’READs [11], PAT-seq [32], TAIL-seq [33, 34], and several others (reviewed in [35–37]). Although these 3’ end sequencing (3’ seq) approaches are powerful and highly sensitive in detecting the precise locations of pAs, even for lowly expressed genes, they are too technically demanding and costly to be widely applied in genomic research. Alternatively, a myriad of computational tools [38–41] have been developed for identifying and quantifying pAs by leveraging the explosively growing RNA sequencing (RNA-seq) data from diverse biological conditions, cell types, individuals, and organisms (Figure 1D). In recent years, the single-cell RNA-seq (scRNA-seq) techniques, particularly those 3’ tag-based protocols such as CEL-seq [42] and 10x Chromium [43], provide great potential to explore dynamics of APA usage during the process of cellular differentiation. Accordingly, a wide spectrum of tools have been proposed to profile APA from diverse scRNA-seq datasets at cell-type or even single-cell resolution [44–46] (Figure 1E).

**Figure 1.**
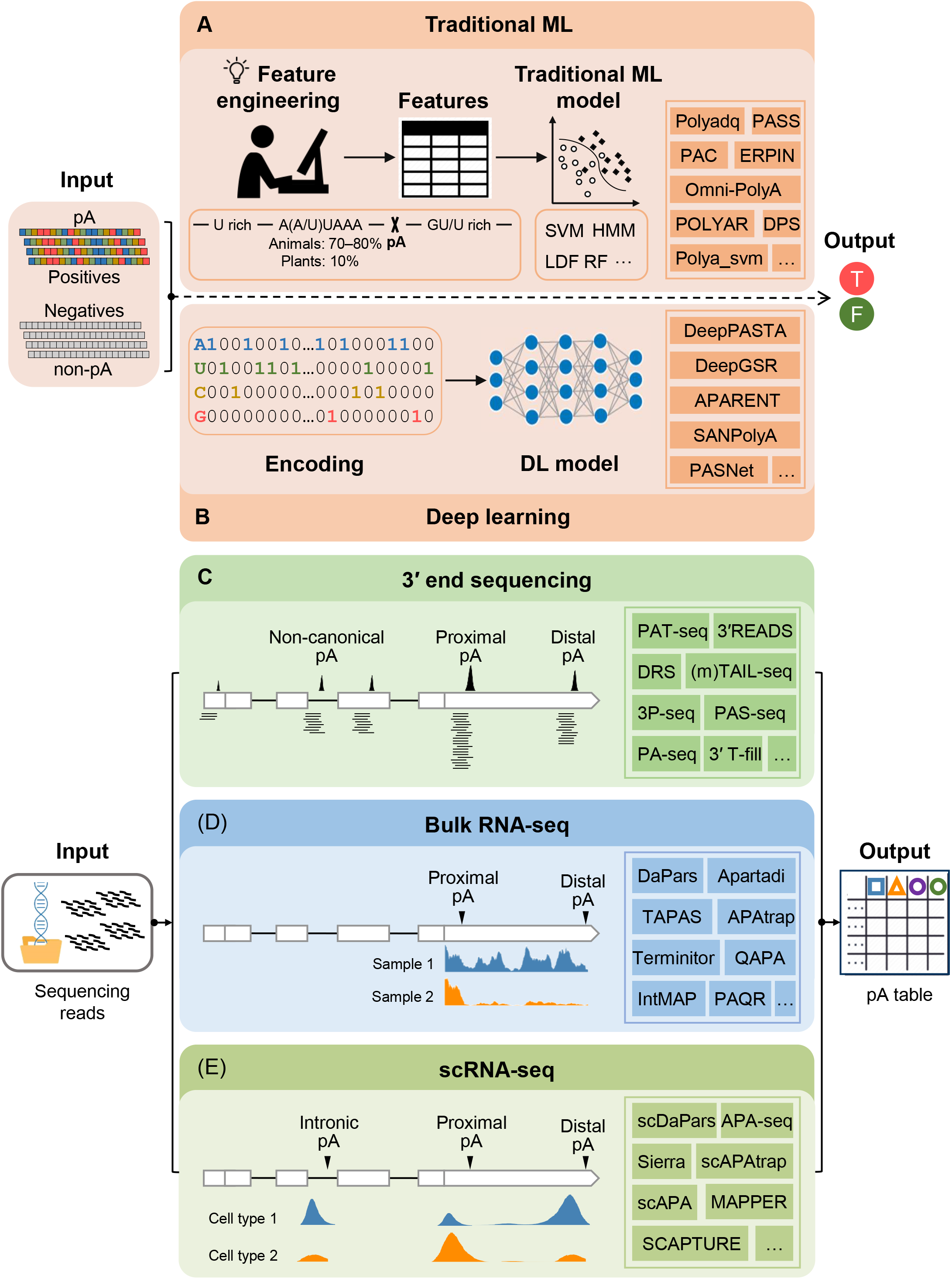
Schematic of computational approaches for predicting pAs from different kinds of sequencing data. **A.** Predicting pAs from DNA sequences based on traditional ML models. **B.** Predicting pAs from DNA sequences based on DL models. **C.** Identifying pAs from 3’ seq data. **D.** Predicting pAs from bulk RNA-seq data. **E.** Predicting pAs from single-cell RNA-seq data. Some representative methods are listed in the text box on the right. pA, poly(A) site; ML, machine learning; T, true; F, false; DL, deep learning; scRNA-seq, single-cell RNA-seq.

The tsunami of genomic data especially bulk and single-cell RNA-seq data and the emergence of ensemble deep learning methodologies have revolutionized computational methods for detecting pAs from diverse kinds of data. In the past decade, a few literature reviews have involved the computational tools for bioinformatic analysis of APA. In 2015, our group summarized computations tools for predicting pAs from DNA sequences and 3’ seq methods for mapping pAs [37]. Szkop and Nobeli [47] described experimental methods for probing 5’ UTRs and 3’ UTRs, and listed computational methods for discovering alternative transcription start sites (TSSs) and pAs from microarray and RNA-seq. Yeh et al. [48] reviewed experimental methods and technologies for studying APA, and briefly listed seven RNA-seq tools for analyzing APA dynamics in tabular form. Chen et al. [49] comprehensively reviewed 3’ seq methods for probing pAs, while their review did not cover the computational tools for APA analysis. Gruber and Zavolan [12] highlighted the importance of APA in health and disease, and briefly listed computational resources for studying APA in a table, including four pA databases, two databases of RBP binding motifs, eight RNA-seq tools for identifying and/or quantifying pAs, and three tools for APA analysis. Our group [50] benchmarked 11 tools for predicting pAs or dynamic APA events from RNA-seq data. Another benchmark study [51] benchmarked five tools for RNA-seq and compared their performance with 3’ seq, Iso-Seq, and PacBio single-molecule full-length RNA-seq method. Ye et al. [52] briefly summarized three computational methods for detecting APA dynamics from diverse single cell types. Zhang et al. [53] focused on the APA regulation in cancer, and briefly listed 14 computational tools for detecting APA. Kandhari et al. [54] highlighted the emerging role of APA as cancer biomarkers and provided an overview of existing relevant experimental and computational methods. However, these two reviews [53, 54] did not distinguish among the prediction of pAs, detection of APA dynamics, and analysis of APA. For example, APAlyzer [55] and movAPA [56] listed in these reviews are actually toolkits for analyzing APA rather than detecting APA dynamics or pAs, which are different from other tools they listed such as DaPars [39] or APAtrap [40]. Generally, although the above reviews have provided detailed overviews of the progress in the complex yet fruitful APA field, none of them has exhaustively summarized available tools for different kinds of data in this field, particularly the emerging DL-based methods and methods for scRNA-seq. Moreover, most reviews only briefly listed tools without delicate summary and sorting, which makes it difficult for the scientific community to decide desirable method for their data analysis. In this review, we described the principles of identifying pAs from different kinds of data and provide an extensive overview of available computational approaches. We catalogued these methods into different categories in terms of the underlying principles of the predictive models and the data they used, and summarized their performance and characteristics such as algorithms, features, and data used in the predictive model. Particularly, we examined several representative tools using RNA-seq and scRNA-seq data from peripheral blood mononuclear cells and put forward operable suggestions on how to assess the reliability of pAs predicted by different tools. We also describe several notes on how to conduct objective benchmark analysis for these massive number of tools. Moreover, we propose practical recommendations on choosing appropriate methods for different scenarios and discussed implications and future directions. Additionally, we highlight outstanding challenges and opportunities using new machine learning and integrative multi-omics techniques. Lastly, we provide our perspective on how computational methodologies might evolve in the future for pA prediction, including non-3’ UTR, tissue-specific, cross-species, and single-cell pA prediction.

## Computational approaches for pA prediction

### Methods for predicting pAs from DNA sequences

The key trigger for cleavage and polyadenylation is the set of *cis*-regulatory elements surrounding a pA, including A[A/U]UAAA hexamer or variant thereof, the UGUA element, upstream and downstream U-rich elements, and downstream GU-rich elements [57]. Since poly(A) signals, the core AAUAAA and its variants, are in the vicinity of most mammalian pAs, the identification of the poly(A) signal (PAS) is usually regarded as an alternative to determine the potential position of a pA. In this review, we refer to the task of predicting pAs or PASs as the “pA identification problem”. During the past few decades, a wide range of computational approaches have been proposed to predict pAs from DNA sequences using experimental and *in silico* mapping of 3’-end expressed sequence tags (ESTs) (Files S1 and S2).

#### Methods based on traditional machine learning models

Earlier studies established traditional ML models to classify a sequence as containing a pA or not, using various algorithms such as discriminant functions [21, 22, 58], hidden Markov model (HMM) [23], SVM [24, 59], Bayesian network [60], artificial neural network and random forests [61], and combined classifiers [25, 62] (**Figure 2** and File S2). The machine learning frameworks of these methods are similar, except that different classification models were employed and/or diverse hand-crafted sequence features were compiled (File S1). As ML models rely heavily on manually designed features and the poly(A) signal of human/animal is considerably different from that of other species like plants or *Saccharomyces cerevisiae* (yeast) [37, 63], these ML-based methods can be divided into two categories according to the applicable species (File S1): i) methods that are applicable to human or animals, including POLYAH [21], Polyadq [58], ERPIN [23], Poly(A) Signal Miner [64], Polya_svm [24], PolyApred [59], POLYAR [22], Chang’s model [65], Dragon PolyA Spotter [61], Xie’s model [66], and Omni-PolyA [25]; ii) methods that are applicable to other species, including the Graber’s method [67] for yeast, POLYA [68] for *Caenorhabditis elegans*, PASS [69, 70], PAC [60] and PASPA [71] for plants, and Wu’s model for *Chlamydomonas Reinhardtii* [62]. These methods utilize diverse sequence features around pAs for pA prediction (File S1). The most commonly used features are position weight matrix for the poly(A) motifs, distance between motifs, and k-gram nucleotide acid patterns [21, 23, 24, 58, 59]. With the increase of the prior knowledge of DNA sequences, more carefully hand-crafted features were derived, such as Z-curve [60], RNA secondary structures [62, 65], physico-chemical, thermodynamic and statistical characteristics [61], the term frequency–inverse document frequency weight [62], and spectral latent features extracted by HMM [66]. Particularly, since the significance of poly(A) signal is different in pAs with different strengths, a few studies divided pAs into sub-groups based on the expression level [22] or pattern assembly [62], and then predicted pAs in each group. In terms of the availability and ease of use of tools, several tools were presented as website (Figure 2), which is particularly convenient for users with little program skill. However, since these tools were generally developed many years ago, the programming languages of many tools are outdated, such as Fortran or Perl, and many tools are no longer available or maintained.

**Figure 2.**
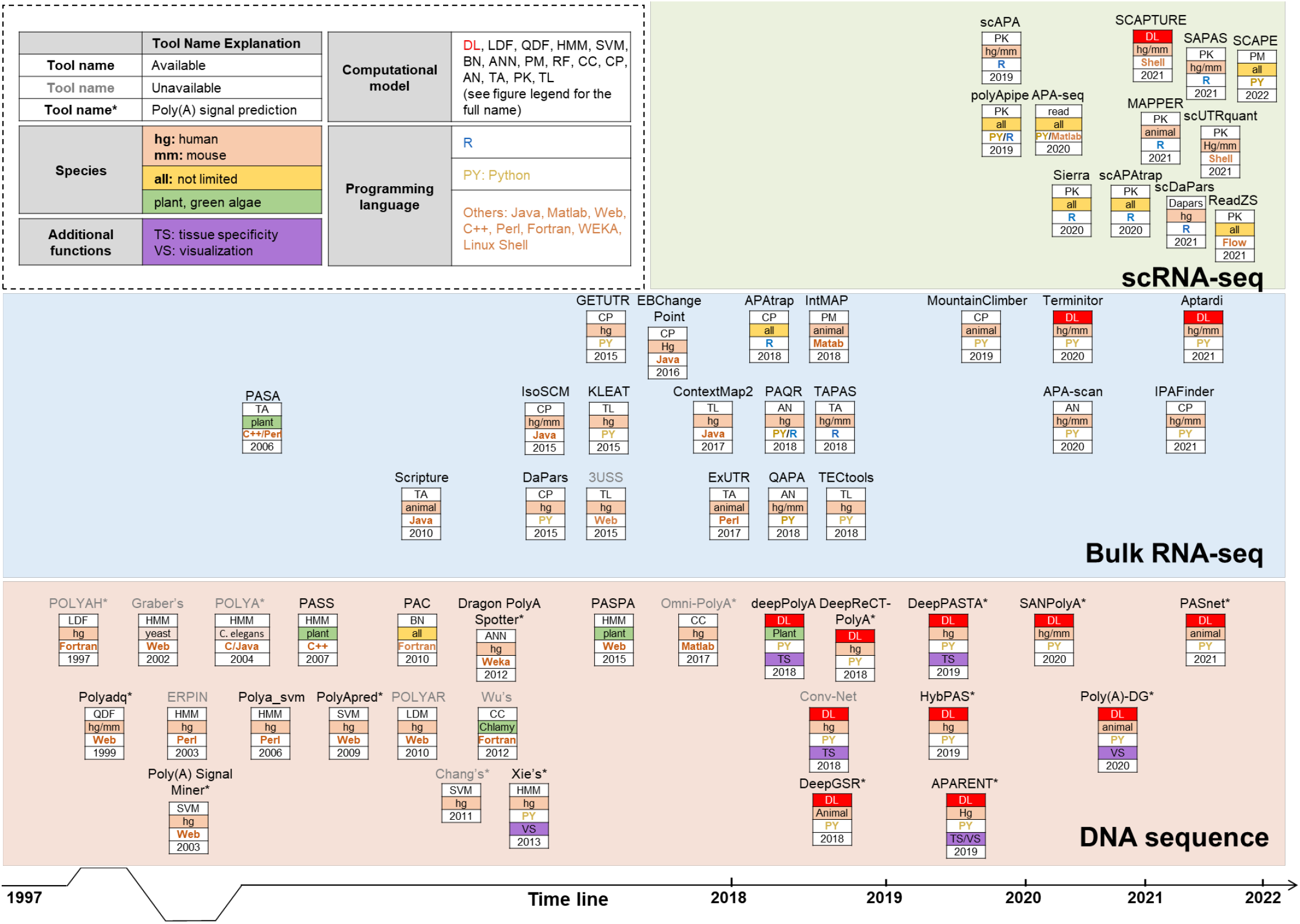
Landscape of computational approaches for predicting pA from DNA sequences, bulk RNA-seq, and single-cell RNA-seq over time. scRNA-seq, single-cell RNA-seq; DL, deep learning; LDF, linear discriminant function; QDF, quadratic discriminant function; HMM, hidden Markov model; SVM, support vector machine; BN, Bayesian network; ANN, artificial neural network; PM, probabilistic model; RF, random forest; CC, combined classifier; CP, change point; AN, annotation-based method; TA, transcript assembly; PK, peak calling; TL, transcript assembly and read linking.

#### Methods based on deep learning models

Despite considerable progress has been made, the overall accuracy and generalizability of traditional ML-based methods remain moderate due to the limited experimentally verified pAs in the early years and the lack of prior domain knowledge to finely design and acquire useful features. In recent years, DL-based methods are emerging rapidly (File S2 and Figure 2), which directly learn hidden features from input nucleotide sequences in a data-driven manner, without knowing any prior knowledge of sequence features. Most methods use convolution neural networks (CNNs), including deepPolyA [72], Conv-Net [73], DeeReCT-PolyA [26], DeepPASTA [28], DeepGSR [27], and APARENT [29]. Other deep learning techniques were also utilized, such as the recurrent neural network (RNN) employed in DeepPASTA [28], a hybrid model with four logistic regression models and eight neural networks used in HybPAS [74], and self-attention mechanisms used in SANPolyA [75] and PASNet [76]. All of these tools were implemented using DL frameworks in Python. In addition to pA prediction, several methods can be utilized for multiple tasks. For example, Conv-Net [73] is capable of inferring pA selection and predicting pathogenicity of polyadenylation variants. DeepPASTA [28] can be used for the prediction of the most dominant pA of a gene in a given tissue and the relative dominance of APA sites in a gene. DeepGSR [27] is able to predict genome-wide and cross-organism genomic signals such as translation initiation sites. APARENT [29] can also be utilized for the quantification of the impact of genetic variants on APA. Different from hand-picked features used in ML-based methods, one-hot encoding features without needing fine feature engineering are widely used in DL-based methods, however, DL-based models are generally of poor interpretability. To enhance the interpretability, several methods provide additional function for visualization of signals. Xia et al. [26] showed the interpretability of their DeeReCT-PolyA model by transforming convolutional filters into sequence logos for the comparison between human and mouse. In APARENT [29], features learned across all network layers were visualized, which can reveal *cis*-regulatory elements known to recruit APA regulators and new sequence determinants of polyadenylation. In addition to performance improvement, DL-based methods have two significant advantages over ML-based methods, the higher generalizability for different species and the higher scalability with large amount of data. For example, DeeReCT-PolyA [26] is an interpretable and transferrable CNN model for recognition of 12 PAS variants, which enables transfer learning across datasets and species. APARENT [29] was trained using isoform expression data from more than three million synthetic APA reporters.

### Methods for predicting pAs from bulk RNA-seq data

Methods that predict pAs only from DNA sequences conspicuously fail to consider *in vivo* expression. RNA-seq has become an indispensable approach for transcriptome profiling in diverse biological samples and a number of methods have been proposed for identifying sample-specific pAs from RNA-seq (File S3). Our group previously benchmarked 11 representative methods for predicting pAs and/or dynamic APA events from RNA-seq [50]. Here we focus on prediction of pAs rather than dynamic APA events. We collected relevant methods summarized in our previous review [50] as well as newly emerging methods, and divided these methods into five categories according to their underlying strategies.

#### Methods that interrogate non-templated poly(A)-capped reads

RNA-seq data contain a small fraction (~0.1%) of non-templated poly(A) tail-containing reads (hereinafter referred to as poly(A) reads) [47], which can be considered as direct evidence for polyadenylation. By interrogating poly(A) reads, an early study [77] identified ~8000 novel pAs in *Drosophila melanogaster* from a total of 1.2 billion RNA-seq reads. Several other methods, such as KLEAT [78] and ContextMap 2 [79], not only employed direct evidence from poly(A) reads but also incorporated transcript assembly to identify pAs. These poly(A) read-based approaches have the advantage to determine the precise locations of pAs, however, it is still challenging to discover pAs of weakly expressed transcripts due to the decreased read coverage near the 3’ end and the low yield of poly(A) reads.

#### Methods based on transcript assembly

Another series of approaches identify pAs from inferred alternative 3’ UTRs by compiling transcript structures from RNA-seq, including PASA [80], Scripture [81], 3USS [82], and ExUTR [83]. These transcriptome assembly-assisted methods deduce gene models first using transcriptome assembly tools, and then identify 3’ UTRs that are absent in the deduced gene models, which rely heavily on assembled gene structures. It is widely accepted that transcriptome assembly from RNA-seq is a rather difficult and computationally demanding task, and it is more challenging to precisely determine 3’ UTRs, especially for lowly expressed genes, due to 3’ biases of read coverage inherent in RNA-seq. Therefore, the performance of these methods is inevitably hindered by potential limitations of existing transcriptome assembly tools.

#### Methods that rely on prior annotations of pAs

During the last decade, numerous experimental techniques have been developed to direct sequence 3’ ends of mRNAs, such as 3’ T-fill [84], 3’READs [11], TAIL-seq [33, 34], to name a few (Figure 1C). Accordingly, several pA databases built upon 3’ seq data of diverse species were continuously released, including PolyA_DB 3 [85], PolyAsite 2.0 [8], and PlantAPAdb [86]. These databases provide a large number high-confidence pAs, which can be used for establishing pA prediction models and evaluating pA prediction results. It is thus naturally to incorporate annotated pAs for predicting pAs from RNA-seq. Several methods, including QAPA [38], PAQR [87], and APA-scan [88], that rely on pre-defined pA annotations were proposed for predicting pAs from RNA-seq. For these methods, the quality of annotated pAs is particularly critical. Most studies establish a comprehensive compendium of well-annotated pAs by merging non-redundant annotations from diverse sources. By combining priori annotated pAs with RNA-seq, the quality of predicted pAs can be greatly improved. However, currently available pA databases are far from complete and limited to only a few well-studied species, such as human, mouse, and *Arabidopsis thaliana*, consequently, these tools are not capable of detecting novel pAs beyond existing poly(A) annotations.

#### Methods that infer pAs by detecting significant changes in RNA-seq read density

Majority of recent approaches predict pAs by modelling read density changes in terminal exons, including GETUTR [89], IsoSCM [90], DaPars/DaPars2 [39, 91, 92], EBChangePoint [93], APAtrap [40], TAPAS [41], moutainClimber [94], and IPAFinder [95]. According to our previous benchmark on 11 tools for RNA-seq [50], TAPAS generally obtained higher sensitivity than other tools across different datasets. Of note, unlike most methods that require at least two samples for change point detection, moutainClimber [94] is a *de novo* cumulative-sum-based approach, which runs on a single RNA-seq sample and simultaneously recognizes multiple TSSs or APA sites in a transcript. Using mountainClimber, Cass and Xiao analyzed 2,342 GTEx samples from 36 tissues of 215 individuals and found 75% of genes exhibited differential APA across tissues [94]. Different from most pA prediction tools focusing mainly on 3’ UTR, IPAFinder was specifically proposed for identifying intronic pAs from RNA-seq [95]. Zhao et al. applied IPAFinder to pan-cancer datasets across six tumor types and discovered 490 recurrent dynamically changed intronic pAs [95]. Methods falling within this category rely on the detection of read density fluctuations which require sufficient read coverage in terminal exons to detect APA sites. It is worth noting that data pre-processing (normalization or smoothing) is particularly important for reducing technical biases caused by non-biological variability [47]. Particularly, some methods, such as APAtrap and DaPars, re-define terminal exon boundaries based on RNA-seq read coverage before identifying pAs, which are capable of detecting pAs in previously unannotated regions.

#### Methods based on machine learning models

In recent years, some newly emerging methods employ traditional ML or DL model to identify pAs from RNA-seq, including TECtools [96], IntMAP [97], Terminitor [98], and Aptardi [99]. TECtools [96] first identifies terminal exons and transcript isoforms ending at known intronic pAs. Then a model was trained based on the aligned RNA-seq data for distinguishing terminal exons from internal exons and background regions, using diverse features reflecting differences in read coverage of these regions. TECtool can also be applied on scRNA-seq, which first pools reads of all cells to infer new transcripts and then quantify each transcript in individual cells. IntMAP [97] leverages one unified ML framework to combine the information from RNA-seq and 3’ seq to quantify different 3’ UTR isoforms using a global optimization strategy. Terminitor [98] is based on a deep neural network for three-label classification problem, which can determine whether an input sequence contains a pA with poly(A) signal, a site without poly(A) signal, or non-pA. Aptardi [99] is a multi-omics approach based on bidirectional long short-term memory recurrent neural network (biLSTM), which predicts pAs by leveraging DNA sequences, RNA-seq, and the predilection of transcriptome assemblers.

### Methods for predicting pAs from single-cell RNA-seq

Single-cell RNA-seq is a powerful high-throughput technique for interrogating transcriptome of individual cells and measuring cell-to-cell variability in transcription [100]. Particularly, several 3’ tag-based scRNA-seq methods enriching for mRNA 3’ ends via poly(A) priming, such as CEL-seq [42], Drop-seq [101], and 10x Chromium [43], provide great potential to dissect APA at single-cell resolution. However, the extremely high dropout rate and cell-to-cell variability inherent in scRNA-seq makes it difficult to directly apply bulk RNA-seq methods to scRNA-seq data. During the last few years, a wide range of computational approaches specifically designed for pA identification from scRNA-seq have emerged (File S4 and Figure 2). We divided these methods into three categories according to their underlying strategies.

#### Methods based on peak calling

The peak calling strategy is widely used by most methods for pA identification from scRNA-seq, including scAPA [102], polyApipe (https://github.com/MonashBioinformaticsPlatform/polyApipe), Sierra [44], scAPAtrap [45], SAPAS [103], and SCAPE [104]. The underlying principle of these methods is that aligned reads from 3’ tag-based scRNA-seq accumulate to form peaks at genomic intervals upstream of pAs [102]. In scAPA [102], a set of non-overlapping 3’ UTRs is first defined from the genome annotation and then peaks within 3’ UTRs are identified using an existing peak calling tool. As adjacent pAs may situate in a single peak, the Gaussian finite mixture model was implemented in scAPA to split large peaks into smaller ones. polyApipe is a pipeline for identifying pAs from 10x Chromium scRNA-seq, which defines peaks of polyA-containing reads. Sierra [44] employed the splice-aware peak calling based on Gaussian curve fitting to determine potential peaks with pAs and then the peaks were annotated and quantified in individual cells. Our group proposed scAPAtrap [45] for identifying and quantifying pAs in individual cells from 3’ tag-based scRNA-seq. scAPAtrap incorporates a genome-wide sensitive peak calling strategy and poly(A) read anchoring, which can accurate locate pAs without using prior genome annotation, even for those with very low read coverage. Yang et al. proposed SAPAS for identifying pAs from poly(A)-containing reads and quantifying pAs in peak regions determined by a parametric clustering algorithm [103]. They further applied SAPAS to the scRNA-seq data of GABAergic neurons and detected cell type-specific APA events and cell-to-cell modality of APA for different GABAergic neuron types. Very recently, Zhou et al. proposed the SCAPE method based on a probabilistic mixture model for identification and quantification of pAs in single cells by utilizing insert size information [104]. The parametric modeling of peaks in most tools based on peak calling such as scAPA or Sierra may cause biases and reduce statistical power in detecting APA events. Alternatively, ReadZS [105], an annotation-free statistical approach, was proposed to characterize read distributions that bypasses parametric peak calling and identify differential APA usages at single-cell resolution among ≥ 2 cell types. ReadZS can not only detect pAs in normal peak shape, but also identify distributional shifts that are not.

#### Methods that rely on prior annotations of pAs

In contrast to the peak calling-based methods used for *de novo* pA identification, a few approaches identify pAs base on prior pA annotations, including MAPPER [106], SCAPTURE [107], and scUTRquant [108]. Li et al. developed MAPPER [106] for predicting pAs from both bulk RNA-seq and scRNA-seq data, which incorporates annotated pAs in PolyA_DB 3 [85] and pools single cells of the same type to mimic pseudo-bulk samples. MAAPER also provides a likelihood-based statistical framework for analyzing APA changes and can identify common and distinct APA events in cell groups from different individuals. The group of MAPPER later developed SCAPTURE [107] which embedded a DL model DeepPASS for evaluating called peaks from scRNA-seq. The DL model was trained by sequences shifting, using annotated pAs from PolyA_DB 3, PolyA-seq, PolyASite 2.0 and GENCODE v39. The authors used SCAPTURE to profile APA dynamics between COVID-19 patients and healthy individuals, and found the preference of proximal pA usage in numerous immune response-associated genes upon SARS-CoV-2 infection. Fansler et al. developed scUTRquant [108] for measuring 3’ UTR isoform expression from scRNA-seq, which relies on a cleavage site atlas established from GENCODE annotation and a mouse Microwell-seq dataset of 400,000 single cells [109].

#### Other methods for predicting pAs from scRNA-seq

Additionally, some other methods do not use the peak calling strategy, including APA-Seq [110] and scDaPars [46]. Levin et al. [110] designed the APA-seq approach to detect and quantify pAs from CEL-seq, which interrogates the gene identity and poly(A) information in the paired Read 1 and Read 2. Although APA-Seq is in principle applicable to other 3’ tag-based scRNA-seq methods, it may not be universally applied in practice in that only sample barcodes rather than the whole 3’ end sequence of the transcript are retained in Read 1 of many public scRNA-seq data [45]. Unlike most tools that are only applicable to 3’ tag-based scRNA-seq, scDaPars [46] that was proposed by the group of DaPars [39] can identify and quantify APA events from either 3’ tag (e.g., 10x Chromium) or full-length (e.g., Smart-seq2) scRNA-seq. In the scDaPars pipeline, DaPars, a tool for identifying APA events from bulk RNA-seq, was first adopted to calculate raw relative APA usage in individual cells, and then a regression model was utilized to impute missing values in the sparse single-cell APA usage matrix. By applying scDaPars to cancer and human endoderm differentiation data, Gao et al. revealed cell type-specific APA regulation and detected novel cell subpopulations that were not found in conventional gene expression analysis.

### Methods for APA analysis rather than pA prediction

In addition to the task of pA prediction (hereinafter termed task 1), there are additional tasks related to the bioinformatic analysis of APA, mainly including the prediction of tissue-specific pAs (task 2), prediction of dominant pAs (task 3), prediction of APA site switching (task 4), and other kinds of APA analysis (task 5). Although most tools described in this review are developed for task 1, several tools are capable of performing multiple tasks. For example, DeepPASTA [28] is able to perform tasks 1-3; Conv-Net [73] can perform tasks 1/3. In this review, we focus only on tools that are applicable to task 1. Of note, NGS-based techniques specially designed for probing pAs, generally known as 3’ seq, such as DRS [10, 30], 3P-Seq [7, 31], and 3’READs [11], are experimental methods rather than computational methods for identifying pAs. Genome-wide pAs generated from 3’ seq are highly confident and are usually regarded as the true reference (i.e., prior information) for building models or evaluating computational methods. These 3’ seq methods are beyond the scope of this review, while have been reviewed in several other reviews [12, 47, 49, 54]. In addition, we have briefly summarized tools or resources designed for APA analysis rather than pA prediction in File S5. Tools such as DeeReCT-APA [111], polyA code [112], and TSAPA [113] are not targeted at task 1 but for other tasks 2/3, such as predicting tissue-specific pAs. Among the five tasks, detection of APA site switching (Task 4) is usually a routine step involved in the analysis of RNA-seq or scRNA-seq. APA site switching reflects the differential usage of APA sites between samples, which does not necessarily need the prediction of pAs (task 1) as a prerequisite. Of note, there are other commonly used phrases similar to ‘APA site switching’ mentioned in this review, such as differential APA site usage [8, 39], 3’ UTR shortening/lengthening [45, 102], and APA dynamics [39, 45, 99, 114]. Some approaches for RNA-seq, such as PHMM [115], ChangePoint [116], MISO [117], and roar [118], directly discover APA site switching by detecting sudden change of read density at terminal exons without identifying APA sites. Recently, several tools were developed for scRNA-seq, such as SCUREL [119], scMAPA [120], and scDAPA [121]. For example, our group developed scDAPA [121] for characterizing differential usages of APA in different cell types using 10x Chromium data, and found APA plays important role in acute myeloid leukemia [114]. Additionally, some toolkits were developed for routine analyses of APA (e.g., annotation and visualization, task 5) using annotated pAs and/or RNA-seq, such as APAlyzer [55] and movAPA [56], while they are not capable of predicting pAs. These diverse tools provide a wide range of complementary resources and opportunities to address the more complex but fruitful field of APA.

## Discussion

### Performance of pA prediction models

At present, there are only a few benchmark studies that systematically evaluate the performance of different tools. Previously, our group benchmarked 11 tools for RNA-seq [50] and found that the sensitivity of some methods varied greatly among different species. For instance, QAPA [38] performs the second best on human data, while it performs the worst on mouse data. APAtrap [40] is the top performer for Arabidopsis data, while TAPAS [41] performs the best on human or mouse data. Recently, Shah et al. [51] benchmarked five tools for RNA-seq against 3’ seq, Iso-Seq, and a full-length RNA-seq method and found that pAs from 3’ seq and Iso-Seq are more reliable than pAs predicted from RNA-seq. They suggested that incorporating the RNA-seq prediction tool QAPA [38] with pA annotations derived from 3’ seq or Iso-Seq can reliably quantify APA dynamics across conditions.

The performance of different tools described in the respective studies was summarized in Files S1 to S4. Generally, for predicting pAs from DNA sequences, DL-based models significantly outperformed ML-based methods and are more suitable for large-scale analysis, owning to the good ability of automatic feature extraction and scalability for big data analysis (Table S2). For example, DeepPASTA [28] has an area under the curve score over 93% in predicting pAs on a DNA sequence dataset, which performed much better than ML-based tools like PolyAR [22] or Dragon PolyA Spotter [61]. APARENT [29], based on deep neural network, was trained on over three million synthetic APA reporter genes, which overcomes inherent size limitations of traditional biological datasets. In contrast, traditional ML-based methods like POLYAR [22] and Omni-PolyA [25] require a considerable amount of prior knowledge and are unable to cope with the rapidly growing data. In terms of the model generalizability, methods for RNA-seq or scRNA-seq are generally applicable to different species if the reference genome and the genome annotation are available. In contrast, the cross-species applicability of methods for DNA sequences is more complex. Models applied to human are normally applicable to other mammals like mouse due to similar poly(A) signals among mammals [57]. However, although most models can be in principle trained using data from a different species, users need collect training data from the other species which are not always available, and most models use hand-crafted features that may not be generalized well across species. Recent techniques like deep learning and transfer learning greatly enhance the generalizability of models. Cross-species experiments have been performed for evaluating the generalizability of some tools, such as DeepGSR [27] and Poly(A)-DG [122], and for these tools single model trained over one species can be generalized well to datasets of other species without retraining. We need to point out that, the evaluation results in a single study may be biased and should be treated with caution, because different datasets and performance indicators were used for the performance evaluation in different studies (Files S1 to S4). In the following section of Conclusions and prospects, we also put forward several notes on how to conduct more objective benchmarking in order to make a fairer comparison of different tools.

### How reliable are the obtained results?

Currently, there is no benchmark evaluation of tools for DNA sequence or scRNA-seq data. Here we attempted to make a preliminary examination of the reliability of results obtained from different pA prediction tools, using a matched bulk RNA-seq and 10x Chromium scRNA-seq data of human peripheral blood mononuclear cells (PBMCs) (File S6). We chose representative tools from each category, including DaPars2 [91], TAPAS [41], and Aptardi [99] for bulk RNA-seq, Sierra [44], scAPAtrap [45], and SCAPTURE [107] for scRNA-seq, and DeepPASTA [28] for DNA sequences (**Figure 3**A top). We collected a total of 676,424 non-redundant pAs from GENCODE v39, PolyASite 2.0, and PolyA_DB 3, which were compiled from 3’ seq and can be used as the true reference (Figure 3A bottom). The number of pAs predicted by different tools, even those under the same category, varies greatly (Figure 3B, left). For example, the number of pAs predicted from RNA-seq by TAPAS and DaPars was nearly 8 times and 4 times that of Aptardi. The number of pAs predicted from scRNA-seq by Sierra and scAPAtrap is about twice that of SCAPTURE. Of note, scAPAtrap can predict pAs for the whole genome including intergenic regions and all the three tools predict a large number of pAs in introns (Figure 3B, right). If only 3’ UTR regions are considered, the number of pAs predicted by the three scRNA-seq tools is much closer (Figure 3C, left). As most tools only identify pAs in 3’ UTR, here we used 3’ UTR pAs for subsequent evaluation. Next, we assessed the authenticity of the predicted pAs by checking whether they are supported by annotated pAs in the true reference. The overlap of pAs predicted from RNA-seq with annotated pAs is much lower than that of scRNA-seq (Figure 3C, left). Particularly, the overlap rate between pAs predicted by SCAPTURE and annotated pAs is as high as 96%, which may be because that the DL model embedded in SCAPTURE was trained with annotated pAs. The position of pAs predicted by TAPAS, SCAPTURE, and scAPAtrap is much more precise than that by other tools (Figure 3C, right). Further, we examined the consistency of the results predicted by different tools. Generally, the consistency among different tools is very low (Figure 3D). For RNA-seq data, only 289 pAs were identified by all the three tools, whereas the vast majority of pAs were identified exclusively by a single tool (Figure 3D, top). In contrast, the consistency of pAs predicted from scRNA-seq by different tools is relatively higher (Figure 3D, bottom). In addition, we assessed the reliability of predicted pAs by investigating sequence features. The single nucleotide profile around pAs predicted by TAPAS, SCAPTURE, and scAPAtrap resembles the general profile [50] (Figure 3E), which is also consistent with the fact that they determine more precise locations for pAs (Figure 3C, right). The percentage of AATAAA around pAs predicted by scRNA-seq is much higher than that of bulk RNA-seq (Figure 3F), indicating that predicted pAs from scRNA-seq tend to be more reliable and more accurate than that from bulk RNA-seq. Next, we used the pA prediction tool for DNA sequence, DeepPASTA, to examine how many pAs identified from RNA-seq data are predicted as true solely based on the sequence characteristics. We extracted the upstream and downstream sequences of pAs predicted by RNA-seq tools as the input for DeepPASTA. The proportion of pAs obtained by different tools to be predicted as true by DeepPASTA is not high and varies greatly, ranging from 28% to 79% (Figure 3G, top), indicating again the low overlap of pAs predicted by different tools. Considering only positive pAs by DeepPASTA, the percentage of AATAAA and 1-nt variants of different tools increased slightly (Figure 3G, bottom vs., Figure 3F), reflecting that positive pAs confirmed by DeepPASTA are relatively more reliable than negative ones. Finally, we examined predicted pAs of the immunoglobulin M heavy chain (*IGMM*) gene, which was reported to express a secreted form using the proximal pA and the membrane-bound form using the distal one [123]. The proximal pA of *IGHM* has been recently found preferentially used in B cells and plasma cells of COVID-19 samples [107]. SCAPTURE and scAPAtrap predicted the precise location of both proximal and distal pAs from scRNA-seq data, while Sierra only predicted the proximal one (Figure 3H). TAPAS predicted three pAs from bulk RNA-seq data, of which two perfectly matched the reference pAs in PolyASite 2.0. In contrast, Aptardi failed to predict any pA for this gene and DaPars2 predicted two pAs yet not verified by reference pAs.

**Figure 3.**
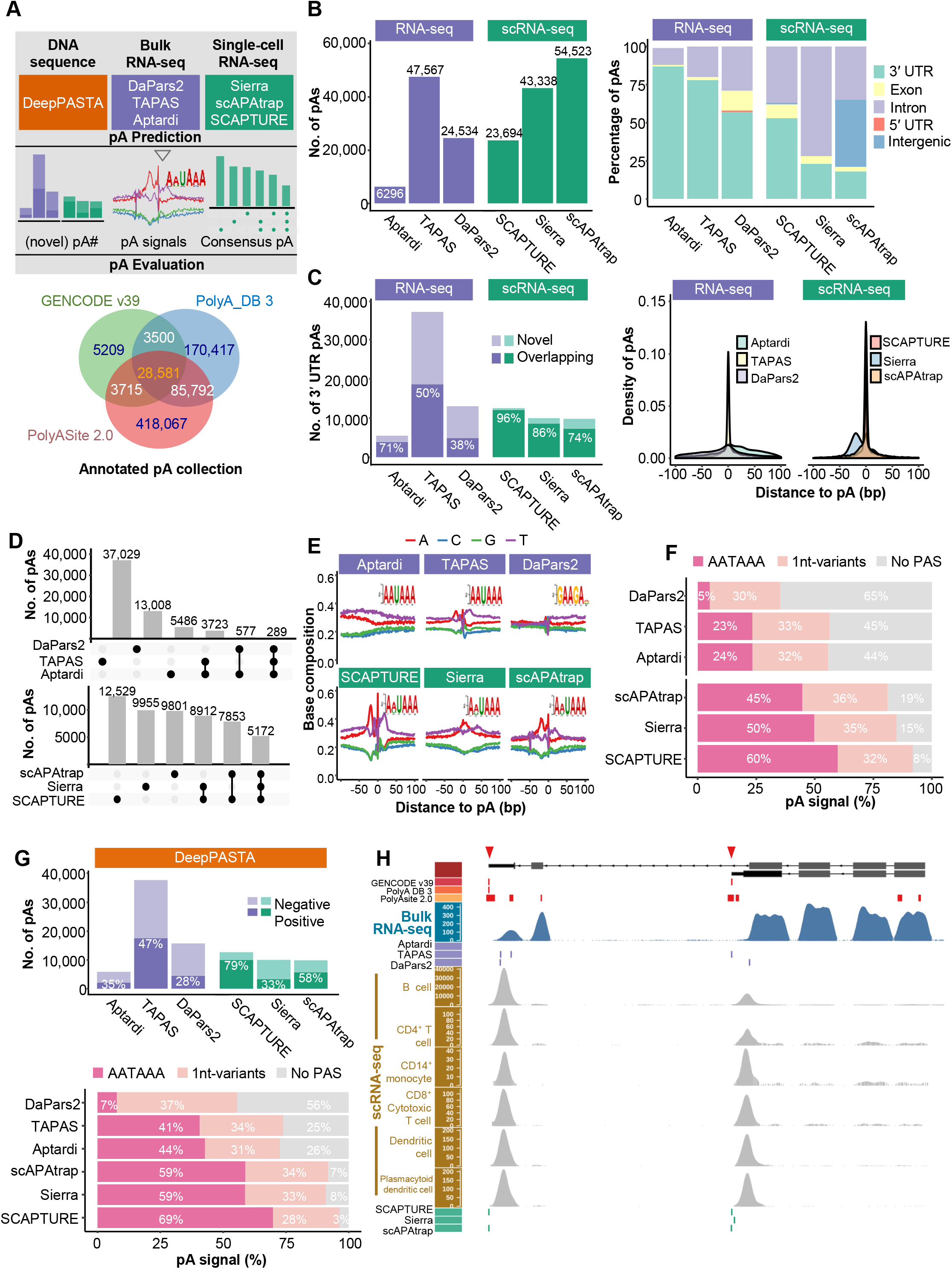
Comparison of representative tools for predicting pAs from a matched bulk RNA-seq and scRNA-seq data of human PBMCs. **A.** Schematic of the benchmark (top) and the collection of reference pAs from GENCODE v39, PolyASite 2.0, and PolyA_DB 3 (bottom). **B.** Number of pAs obtained by different tools (left) and distributions of pAs in different genomic regions (right). **C.** Overlap of 3’ UTR pAs predicted by different tools with reference pAs (left) and distributions of distance from predicted 3’ UTR pAs to reference pAs (right). **D.** Overlap of 3’ UTR pAs predicted by different tools from RNA-seq data (top) and scRNA-seq data (bottom). **E.** Single nucleotide profile around 3’ UTR pAs predicted by different tools. For each tool, the sequence logo of the most dominant motif around the pA identified by DREME was also shown. **F.** Number of occurrences of AATAAA and 1-nt variants around pAs predicted by different tools. **G.** The proportion of pAs obtained by different tools to be predicted as positive or negative by DeepPASTA (top) and the number of occurrences of AATAAA and 1-nt variants around positive pAs (bottom). The upstream and downstream sequences of pAs predicted by each RNA-seq tool were extracted as the input for DeepPASTA. **H.** Predicted 3’ UTR pAs by different tools for the *IGHM* gene. Tracks from top to the bottom are gene model, reference pAs from three databases, read coverage from bulk RNA-seq, predicted pAs from bulk RNA-seq, read coverage for each cell type of scRNA-seq, and predicted pAs from scRNA-seq. The red triangles on the chromosome strip highlight the two representative pAs of *IGHM*.

Although this preliminary benchmark is far from objective or exhaustive to reflect the advantages and disadvantages of different tools, it reveals several potential issues when using the results obtained by different pA prediction methods. First, although a considerable number of pAs were identified by most tools, the overall prediction accuracy and sensitivity of these tools is low (Figure 3C). Our previous comparative study [50] on tools for bulk RNA-seq have also revealed that a considerable number of predicted pAs were not annotated in 3’ seq, and the overall prediction accuracy of these tools, even the best one, TAPAS, is not high (40%–60% for human/mouse data). It is still challenging to determine whether a pA not present in prior annotations is false or novel. We anticipate that at least part of predicted pAs that are not overlapping with annotated ones may potentially be true due to that the current pA annotations are still far from complete. Second, the number of pAs identified by different tools, either for bulk RNA-seq or scRNA-seq, varies greatly, and the consensus of results obtained by different tools is limited (Figure 3D). This is also similar to the observation in our previous benchmark that each tool predicts an independent set of pAs and the overlap of results from different tools is extremely low (< 7% for human/mouse data) [50]. Third, as some tools incorporate additional information to predict pAs, e.g., prior pAs used by SCAPTURE and poly(A) reads used by scAPAtrap, the resolution of pAs predicted by different tools varies greatly (Figures 3C&E). Fourth, 21% to 72% of the predicted pAs by different tools were not recognized as true pAs based on their sequence features (Figure 3G). Fifth, although scRNA-seq data suffers from extremely high level of noise and sparsity, prediction results from scRNA-seq seem to be more reliable and consistent than those from bulk RNA-seq (Figures 3C, D &F). However, this is not unexpected because that it may be less challenging to computationally predict pAs from the 3’-tag based scRNA-seq data than the full-length-based bulk RNA-seq data. Still, further benchmark study with more complete prior annotations, diverse datasets, and performance indicators is needed in order to assess the results obtained from different tools more fairly and objectively.

Here we try to give some operable suggestions on how to obtain high-confidence pAs. The most straightforward way may be making a consensus set of pAs that are predicted by multiple tools, however, this may result in a relatively small number of pAs due to the limited overlap by different tools. Another way is to obtain the intersection of predicted sites and real sites, using annotated pAs that are manually curated and available in several databases such as PolyASite 2.0 and PolyA_DB 3. However, it should be noted that these annotated data sources were compiled from limited biological samples and species; they are far from complete to cover all real sites especially tissue-specific ones. Similar to our benchmark analysis on RNA-seq and scRNA-seq PBMCs (Figure 3), users can also use data from another omics from similar biological samples, if available, to predict pAs for mutual verification. In addition, since many sequence motifs, e.g., AAUAAA and its variants, have been reported to have a positional preference relative to the pA, it is naturally to examine sequence patterns surrounding each predicted pA to get pAs with explicit poly(A) signals. This is particularly useful for assessing the authenticity of pAs from animals because AAUAAA and its 1-nt variants appeared in > 90% of animal pAs [8]. In contrast, AAUAAA only accounts for < 10% of pAs in plants, therefore it is not practical to validate plant pAs through sequence features. Moreover, the general single nucleoside compositions surrounding pAs in different species have been clearly reported, we can thus inspect the base composition around predicted pAs. Of note, this way is applicable to evaluation of the overall quality of the pAs, while it cannot be used to assess the reliability of a single pA. The movAPA package [56] can be used for most of the above-mentioned quality assessments.

### Practical guidelines for choosing appropriate methods

Based on the summary of different methods (Files S1-S4), we attempt to choose representative tools from each category and propose a set of practical guidelines for users (**Table 1**). As methods in different categories use different kinds of data as the input, the choice of the method first depends on the users’ own data. For bulk RNA-seq data, the choice of the method should be mainly driven by the availability of pA annotations. For scRNA-seq data, the choice of the method mainly depends on the protocol of the scRNA-seq (e.g., 3’ tag or full-length) and the availability of pA annotations. For methods predicting pAs from DNA sequence, the choice of the method should be primarily driven by the algorithm used, deep learning or traditional machine learning. Particularly, for cross-species pA prediction from DNA sequences, users should pay extra attention to whether they need to retrain the model for individual species, which may require users to have certain programming ability. Additionally, several tools are in the form of web servers, providing a portable platform for predicting pAs from DNA sequences for researchers with limited programming ability. Several other factors also affect the choice of methods, such as the availability of the tool or code, the popularity, the ease of use, the clarity of documentation, and the scale of the data. When predicting pAs on a dataset of interest, it is important to further consider two points. First, it is critical that the obtained pAs and/or the downstream results (e.g., differential APA events) are confirmed by multiple pA prediction methods. This is to ensure that the prediction is not biased due to predefined parameter settings or the specific algorithm used in the method. The merit of using different methods is also demonstrated by the benchmark results in previous studies [50, 51] and in this study (Figure 3), which show substantial complementarity between different methods. Second, even if prior pA annotations are available, it can be also beneficial to try out methods that do not rely on prior annotations. When predicted pAs, even a small portion, are confirmed using such a different method, it provides users with additional evidence.

## Conclusions and prospects

### Challenges in improving the performance of pA prediction

The field of pA prediction is progressing rapidly, primarily in the aspects of using DL models and predicting pAs at the single-cell resolution. However, the overall accuracy, sensitivity, and specificity of currently available methods remain moderate (Figure 3). The coming flood of extensive sequencing data, especially multi-omics and single-cell data, will provide new opportunities but also demand new computational methods to exploit this new information. Potential challenges of improving the prediction performance include but are not limited to: paucity of annotated pAs covering diverse tissues and species; mis-assemblies caused by the low complexity 3’ UTR sequences; mis-alignment of short reads or incomplete sequence coverage near 3’ ends; difficulty in capturing pAs in low-expression genes; poor knowledge on primary, secondary or higher structure information of poly(A) signals, particularly in plants; gaps in our knowledge on understanding APA regulators in different omics layers; limited success in integrating the quantitative features from multiple omics layers; lack of transferrable intelligent methods for cross-species prediction; lack of interpretability in models based on deep neural networks; hurdle in constructing negative datasets due to the prevalence of unconventional pAs in CDS and introns; difficulty in identifying multiple pAs anywhere in the transcript; lack of effective algorithms to deal with the extremely high isoform-level dropout rate and noise inherent in scRNA-seq. Furthermore, higher standards for software quality assurance and documentation would help improve the ease of use of these tools and facilitate their application in the broader community. Finally, new algorithms should be designed to cope with ever-increasing amount of different kinds of data, especially the explosion in single-cell data with multi-omics features.

### Notes on benchmarking different methods for predicting pAs

Till now, there are few reports on the exhaustive evaluation of computational tools for predicting pAs. Previously, our group benchmarked 11 representative tools for predicting pAs and/or dynamic APA events from RNA-seq [50]. Lately, Shah et al. [51] evaluated five tools for RNA-seq against 3’ seq, Iso-Seq, and a full-length RNA-seq method in identifying pAs and quantifying pA usage. However, there is no study to provide an exhaustive evaluation of existing tools for pA prediction from different kinds of data, particularly those tools for scRNA-seq. Here we attempt to give some notes on benchmarking analysis in this field. First, the real pA dataset is very critical for performance evaluation, however, the reference datasets used in different studies are quite different. Therefore, it is imperative to compile reliable reference datasets with uniform standards. In particular, RNA-seq or scRNA-seq data are sample-specific, so the reference pA dataset from matched samples should also be considered. Moreover, due to the paucity of real pA dataset at the single-cell level, possible deviations need to be considered when using real pA data from bulk data for evaluation. For example, pAs exclusively recognized in single cells may be authentic pAs from rare transcripts or rare cells, even though they may not be present in the bulk pA reference. Second, most tools were evaluated using data only in mammals (mainly human and mouse), therefore the scalability of these tools in different species, especially their applicability to plants, needs to be further evaluated. Third, almost all published prediction tools provide their own benchmark pipelines using different datasets, which potentially favors their prediction efficiency. These benchmark protocols might be credible, but may lack objectivity, simplicity, and effectiveness. We have sorted out the data used for performance evaluation in the respective study of each tool in detail (Files S1-S4), which can facilitate researchers to compile more diverse and standard data for objective benchmark in the future. So far, the most widely used datasets for evaluating pA tools for DNA sequences are the PASS dataset [69, 70] of plant species, the ERPIN dataset [23], and DeepGSR dataset [27] of animal species; datasets for RNA-seq are the MAQC dataset [124] and the HEK293 dataset [125]; datasets for scRNA-seq are the 10x human PBMC data and the *Tabula Muris* atlas [126] (Files S1-S4). Moreover, genomic data could be small sample data and large-scale data, it is also necessary to evaluate the performance of different tools under different sizes of data. Fourth, the output format varies among different tools. For example, most tools for DNA sequences generate binary output or probabilities between 0 and 1; some tools for RNA-seq or scRNA-seq output potential regions of pA instead of exact pA position. Therefore, how to unify the output of different tools for objective evaluation needs to be carefully considered. Fifth, compared with the benchmark of tools for DNA sequence data, the benchmark for scRNA-seq tools is much less uniform (Files S1-S4). Almost all studies examined the consensus between the identified pAs and annotated pAs, while there is still no commonly used objective evaluation strategies with diverse indicators. Therefore, it is necessary to use a variety of performance indicators (e.g., sensitivity, specificity, and precision) that are complementary in nature for comprehensive performance evaluation, particularly for the evaluation of the emerging scRNA-seq tools. At the same time, it is also important to simply present an overall ranking of different tools. The last but not least, many tools have parameters that can be adjusted, however, only the default parameters were normally used for evaluation. Therefore, some strategies (e.g., grid search) should be proposed to evaluate the impact of different parameters of a method.

### Predicting pAs in non-3’ UTRs

With the advance of 3’ seq, more and more unconventional pAs located in non-3’ UTR regions like intron and CDS were discovered [3, 49, 127]. These non-3’ UTR pAs may generate mRNA isoforms encoding distinct proteins or result in the creation of premature stop codons. Intronic polyadenylation has been found associated with cancer through the inactivation of tumor-suppressor genes [95, 128]. The differential use of intronic pAs is a potential indicator for the differential expression of pre-spliced mRNA transcripts, which contributes to detecting newly transcribed genes and ultimately helps estimate the rate and direction of cell differentiation [129]. Till now, almost all computational tools focused on pA prediction in 3’ UTRs. Many tools, particularly those for DNA sequences, usually consider random sequences from introns as negative datasets for model training, which would cause some real intronic pAs to be mistakenly regarded as negative instances. Therefore, even for the pA prediction in 3’ UTRs, it is necessary to consider the prevalence of unconventional pAs when constructing the negative dataset. Lately, some tools for bulk or single-cell RNA-seq have found a considerable number of pAs in introns. By applying IPAFinder [95] on pan-cancer data from bulk RNA-seq, 490 recurrent dynamically changed intronic pAs were found. Sierra [44] utilized a splice-aware strategy and identified a considerable number of intronic peaks from scRNA-seq, however, the majority of these peaks may be internal priming artifacts as they are proximal to A-rich regions. SCAPTURE [107] also found > 16,000 candidate intronic pAs from 10x PBMC samples, while < 20% pAs were overlapped with known intronic sites and a large number of false positives were present in lowly expressed genes. Therefore, further careful inspection or filtering is critical to obtain true non-3’ UTR pAs or new intelligent algorithms are demanded to effectively call non-3’ UTR pAs.

### Predicting tissue-specific pAs

APA plays a significant role in tissue-specific regulation of gene expression [2, 12]. Profiling APA dynamics or differential APA usages under different physiological or pathological conditions has become a routine analysis in most APA studies. Computational prediction of tissue-specific pAs may be an alternative yet cost-effective solution for analyzing tissue-specificity of APA. The pA prediction problem described in this review is essentially a binary classification problem, which aims to distinguish between nucleotide sequences or genomic regions that contain a pA and those do not. Studies are currently in progress to solve the problem of pA quantification, which aims to predict the strength or dominance of a given pA across tissues. Weng et al. [112] and Hafez et al. [130] predicted whether a given pA is tissue-specific or not, whereas they do not tackle the question of alternative choice of APA sites. One way to study tissue specificity of pAs is to explore the differential usage of APA sites in a gene (e.g., proximal and distal pAs). Several tools, such as Conv-Net [73], have been proposed to predict the strength of APAs sites. Leung et al [73] predicted relative dominance of pAs within 3’ UTR in human tissues solely based on nucleotide sequences using a DL model. However, these methods only make predictions based on sequence features, while fail to consider sample-specificity and *in vivo* expression. In contrast, many tools for RNA-seq or scRNA-seq can be used for pairwise comparisons between two samples, while they are not very suitable for profiling APA across multiple tissues. Ever-larger RNA-seq or scRNA-seq data comprising of growing number of samples from diverse tissues are increasingly available, which places new demands on developing new methods to efficiently tackle the question of tissue-specificity of APA.

### Cross-species prediction of pAs

Traditional ML methods, such as those based on SVM, can hardly adapt to different species, because they used hand-crafted features learnt for a specific species. Although many DL-based tools have been proposed to improve the performance of pA prediction, most tools still need species-specific real pA collection for model training. Consequently, these tools may suffer from high risks of overfitting and are not applicable to species without any prior pA annotations. Therefore, it is a promising direction to design new transferrable algorithms for cross-species pA prediction or to improve the generalizability of existing tools, which allows a well-trained model from a species with rich annotations to be transferred to data from a different species without retraining or prior knowledge. Annotation-assisted methods, compared to methods without using prior annotations, generally ensure higher data quality and achieve better performance, however their application is limited to data from specific species or biological conditions. Collection of more extensive pA annotations from different sources would definitely contribute to predicting novel sites and increasing the coverage of pAs in diverse cell types, biological conditions, and species. Therefore, an alternative solution for predicting pAs in poorly annotated species could be building an elegant model for well-annotated species and then transferring the model to a different but related species, even without an established pA collection. An initial attempt has been made by some existing methods like Poly(A)-DG [122], which extracts shared features from multiple species and can be generalized to the target species without fine-tuning. However, Poly(A)-DG was only tested between four animals. Till now, tools applicable to plants are still limited. It is widely accepted that the sequence conservation in poly(A) signals in plants is very low, where the most dominant AAUAAA only appears in less than 10% of pAs [57]. Our group recently developed a tool called QuantifyPoly(A) [63] to profile genome-wide polyadenylation choices, which found plant pAs generally exhibit higher micro-heterogeneity than animal ones, and UGUA, UAAA and/or AAUAAA are used in a species-dependent manner. Still, more efforts are needed to explore additional motifs and/or higher-order structures associated with plant polyadenylation and more intelligent algorithms are demanded in order to better predict pAs in multiple species.

### Predicting pAs by integrating multi-omics data

Poly(A) sites can be derived from different kinds of data. For example, 3’ seq has the unique advantage of acquiring high-quality pAs transcriptome-wide, which contributes to a larger compendium of authentic pAs. Third-generation sequencing technologies, such as PacBio sequencing, are powerful in profiling full-length transcriptome, which could provide a more accurate transcriptome annotation. Widely conducted bulk RNA-seq data can be used for capture and quantify pAs of low-abundance transcripts, and the rapid growing scRNA-seq data support the identification of relatively rare transcripts in single cells. In addition to the genome or transcriptome layers, APA modulation has been found associated with other layers of gene regulation, such as nucleosome positioning, transcription rate, DNA methylation, and RNA-binding proteins [2, 131–133]. By integrating multi-omics data, weak signals from one layer can be amplified or noises be reduced to avoid false negative predictions by referring to the complementary information from additional layers. For instance, potential pAs identified from RNA-seq without well-recognized poly(A) signals could be eliminated if there is no evidence in 3’ seq or full-length RNA-seq data. Initial attempts have been made for APA analysis using multi-omics data. scUTRquant [108] incorporates a cleavage site atlas established from a mouse full-length Microwell-seq dataset of 400,000 single cells [109] for filtering high-confidence pAs predicted from 3’ tag scRNA-seq. Leung, et al. [73] predicted strength of pAs using nucleotide sequences, considering features from additional layers like nucleosome positioning and RNA/binding protein motifs. IntMAP [97] is a unified ML-based framework, which can fine-tune the contributions of RNA-seq and 3’ seq data by tailoring the parameter *λ*. Currently, DL models have been widely used in predicting pAs from DNA sequences. However, in many cases, DL models fail to make accurate prediction, while patterns of RNA-seq coverage provide clear evidence of polyadenylation, and vice versa [111]. Accordingly, several DL-based tools integrated bulk or single-cell RNA-seq with DNA sequences for pA prediction, such as SCAPTURE [107] and Aptardi [99]. It is promising yet challenging to formulate one unified computational framework, especially leveraging the strength of intelligent algorithms, to integrate the quantitative information from multiple omics layers, e.g., genomic DNA, transcriptome data, methylation data, and chromatin accessibility data, to identify and quantify pAs genome-wide.

### Predicting pAs at the single-cell level

With the rapid development of scRNA-seq technology, different tools continue to emerge for pA identification in single cells. Currently, most methods, like scAPA [102], Sierra [44], and MAPPER [106], construct pseudo-bulk RNA-seq data by pooling reads from cells of the same cell cluster (or cell type) to address the high dropout rate and variability inherent in scRNA-seq. Although many of these tools, like scAPA or Sierra, can still quantify the expression of a pA in each cell by counting reads within a poly(A) region, single cell-based quantification may have high noise level and missing values due to biological and technical variance [106]. As such, APA usage is characterized at the cell-cluster resolution rather than the single-cell resolution, which somewhat contradicts the ultimate goal of single-cell sequencing. Moreover, cell clusters or cell types in these studies were inferred by the conventional gene-cell expression profile, consequently, the APA analysis is limited to predefined cell types and the result may be affected by different cell type annotations. Alternatively, scDaPars [46] quantifies single-cell APA usage based on the model for bulk RNA-seq introduced in DaPars [39], and then recovers missing APA usage by leveraging APA information of the same gene in similar cells. Another limitation of most tools for scRNA-seq is that they are only applicable to 3’ tag based scRNA-seq like 10x Chromium or CEL-seq. Till now, only scDaPars can be applied to both 3’ tag and full-length scRNA-seq, e.g., Smart-seq2. However, although scDaPars is reported to be able to quantify APA usage in individual cells independent of gene expression, pAs were actually predicted from the bulk RNA-seq tool DaPars that was not specifically designed for scRNA-seq. Moreover, it is challenging to identify and verify low-expression pAs in highly sparse scRNA-seq, particularly those in rare cells. In addition, the tsunami of complex scRNA-seq datasets with various tissue sources, batch effects, and library sizes also have brought huge computational and analytical challenges. Therefore, more efforts are needed to develop new methods to address inherent issues in scRNA-seq for establishing a more comprehensive landscape of pAs at the single-gene and single-cell resolution.

## CRediT author statement

**Wenbin Ye:** Investigation, Data curatin, Visualization, Writing - review & editing. **Qiwei Lian:** Data curation, Writing - review & editing. **Congting Ye:** Investigation, Writing - review & editing. **Xiaohui Wu:** Conceptualization, Writing - original draft, Writing - review & editing, Supervision, Project administration, Funding acquisition. All authors read and approved the final manuscript.

## Acknowledgments

This work was supported by the National Natural Science Foundation of China (Grant No. 61871463 to XW) and the Natural Science Foundation of Fujian Province of China (Grant No. 2020J01047 to CY).

## Tables

**Table 1 Recommended tools for predicting pAs from DNA sequences, bulk RNA-seq, and single-cell RNA-seq**

## Supplementary material

**File S1 List of methods for predicting poly(A) sites or poly(A) signals from DNA sequences based on traditional machine learning models**

**File S2 List of methods for predicting poly(A) sites or poly(A) signals from DNA sequences based on deep learning models**

**File S3 List of methods for predicting poly(A) sites from RNA-seq**

**File S4 List of methods for predicting poly(A) sites from single-cell RNA-seq File S5 List of methods or resources for analysis of alternative polyadenylation rather than prediction of poly(A) sites**

**File S6 Materials and methods used in this study**

