## Supplementary material for "A Survey on Methods for Predicting Polyadenylation Sites from DNA Sequences, Bulk RNA-seq, and Single-cell RNA-seq": File S1

### File S1. Methods for predicting poly(A) sites or poly(A) signals from DNA sequences based on traditional machine learning models.

| | Method | Model | PAS/pA | Site# | Species | Range | Data[1] | Features | Performance | Code | Availability |
|---|---|---|---|---|---|---|---|---|---|---|---|
| ***Tools applicable to species other than human or mouse*** | | | | | | | | | | | |
| 1 | **(Graber, et al., 2002)** | Hidden Markov model (HMM) | pA | Multi | Yeast | –110, +40 | 1) A dataset from (Graber, et al., 1999) with 1352 pAs from ESTs. | A HMM-based probabilistic model to include information from a relative position of –110 to +40 with respect to the 3' processing site (including efficiency, positioning, near-upstream, near-downstream element and pA) | Competing tools: -<br>Indicators: SN<br>Results: SN=0.33, 0.58, 0.67 and 0.87 (allowing a mismatch of 0, 1, 3, 10 nt). | Web server | - |
| 2 | **POLYA (Hajarnavis, et al., 2004)** | Hidden Markov model | AATAAA | 1 | C. elegans | - | 1) 961 sequences with unique AATAAA and cleavage site collected and processed from WormBase. | Six distinct regions in the vicinity of the 3' end modeled by HMM. | Competing tools: -<br>Indicators: SN, SP<br>Results: SN=70.4%, SP=70.4% based on Viterbi-HMM (3' UTR region); SN=81.6%, SP=67.7% based on posterior decoding-HMM (3' UTR region). | C, Java | - |
| 3 | **PASS (Ji, et al., 2007; Shen, et al., 2008)** | Generalized hidden Markov model | pA | Multi | Arabidopsis, rice | -300, + 100 | **[PASS dataset]:**<br>1) 8K dataset: Arabidopsis 3' UTR of 8160 pAs.<br>2) 55K dataset: rice: 55,000 annotated pAs in rice.<br>3) Negative dataset: randomly generated 3' UTRs but preserved the original trinucleotide distribution, 5' UTRs, introns and coding sequences. | The base output probabilities (ATCG), assigning weights for conserved signal patterns (WGH), constructing the first-order heterogeneous Markov sub-model (PNMATRIX), and using their combination (WGHPN). | Competing tools: -<br>Indicators: SN, SP<br>Results: Arabidopsis: SN, SP=97% (coding sequences and random datasets, at threshold=4); SN, SP=72% to 82% (5' UTR and introns, at threshold=5.2 to 6);<br>Rice: best combination of SP and SN (90%) when the threshold was set at 4. | C++ | http://www.bmibig.cn/mnt/paspa/ |
| 4 | **PAC (Ji, et al., 2010)** | Bayesian network | pA | 1 | Arabidopsis, human | -130, +30 | 1) The PASS dataset (Ji, et al., 2007; Shen, et al., 2008). | K-gram pattern, Z-curve, position-specific scoring matrix and first-order inhomogeneous Markov sub-model. | Competing tools: PASS<br>Indicators: SN, SP<br>Results: SN, SP=95% (best combinations, in Arabidopsis) | Delphi | http://www.bmibig.cn/mnt/tools/PAC/ |
| 5 | **(Wu, et al., 2012)** | Combined classifier | pA | 1 | Chlamy | -150, +30 | 1) Positive dataset contains 16,952 sequences with pAs.<br>2) Four negative datasets consist of four types of control sequences without any pAs, including 1,658 5' UTR sequences, 17,032 intron sequences, 11,471 CDS, and 10,000 randomly generated sequences. | Predicted RNA secondary structure, TF-IDF weight, first-order Markov chain, pentamer ratio and a position weight matrix. | Competing tools: -<br>Indicators: SN, SP<br>Results: SN, SP= 83-99% | Delphi | - |
| 6 | **PASPA (Ji, et al., 2015)** | GHMM | pA | Multi | Plants | -300, 100 | 1) **[PASS dataset]** and *M. truncatula*, and *C. reinhardtii* dataset (Ji, et al., 2007; Shen, et al., 2008; Wu, et al., 2012; Zhao, et al., 2014).<br>2) pAs for other six species were collected by processing ESTs downloaded from NCBI. | As in (Ji, et al., 2007). | Competing tools: -<br>Indicators: SN, SP<br>Results: SN and SP in a range between 0.80 and 0.95. | Web server | The website is no longer available, but users can download the tool from http://www.bmibig.cn/mnt/tools/paspa/. |
| ***Tools applicable to human or mouse*** | | | | | | | | | | | |
| 7 | **POLYAH (Salamov and Solovyev, 1997)** | Linear discriminant function | AATAAA | 1 | Human | -100, +200 | 1) 248 positive and 5702 negative pAs in the training set.<br>2) 131 and 1466 in the test set (GenBank, Version 82 ). | Position weight matrix for PAS and downstream element (DE), distance between PAS and DE, hexanucleotide composition, positional triplet composition, positional triplet composition. | Competing tools: -<br>Indicators: SN, SP, CC<br>Results: This tool predicts 86% of poly(A) regions correctly, with a specificity of 51% and correlation coefficient of 0.62. | Fortran | - |

| | | | | | | | | | | | |
|---|---|---|---|---|---|---|---|---|---|---|---|
| 8 | **Polyadq (Tabaska and Zhang, 1999)** | Quadratic discriminant function | AATAAA, ATTAAA | 1 | Human, mouse | -6, +100 | 1) Known pAs contains 280 mRNA sequences and 136 DNA 2) sequences.<br>2) The negative training set contains 462 pseudo signals. | PAS and DE weight matrices. | Competing tools: Quant2, GRAIL, POLYAH<br>Indicators: SN, SP, CC<br>Results: a correlation coefficient of 0.413 on whole genes and 0.512 in the last two exons of genes. | Web server | http://rulai.cshl.org/tools/polyadq/polyadq_form.html |
| 9 | **ERPIN (Legendre and Gautheret, 2003)** | Probabilistic hidden Markov model | pA | 1 | Human | -300, +300 | **[ERPIN dataset]:**<br>1) 4956 EST-validated pAs.<br>2) Control sequences not containing pAs: coding sequences (CDS), introns, and two types of randomized 3' UTR sequence.<br>3) Strong proximal sites: 129; strong distal sites: 499; weak proximal sites: 655; weak distal sites: 210. | Position-dependent compositions in the terminal sequences were measured in an 11-nucleotide sliding window. | Competing tools: polyadq<br>Indicators: SN, SP, ACC<br>Results:<br>A specificity improvement of 9.7% relative to the polyadq method;<br>SN=56%, SP=69.49-85.38% across negative set CDS, introns, UTR shuffled, UTR Markov 1$^{st}$ order. | Perl | - |
| 10 | **Poly(A) Signal Miner (Liu, et al., 2003)** | SVM | NNTANA | <=5 | Human | -100, +100 | **1) [ERPIN dataset]**<br>2) **[PSM dataset]:** 312 human mRNA sequences from RefSeq and negative dataset was generated by scanning for the occurrences of AATAAA at coding region. | K-gram nucleotide acid or amino acid patterns. | Competing tools: polyadq, ERPIN<br>Indicators: SN, SP<br>Results:<br>ERPIN SN=55.9%, Polyadq=55.7%, Poly(A) Signal Miner=56.3%;<br>the sensitivity and specificity of 10-fold cross-validation on training data are 89.3% and 80.5%;<br>for the ERPIN data, specificity of 73% to 93% at the same sensitivity (56%). | Web server | http://dnafsminer.bic.nus.edu.sg/PolyA.html |
| 11 | **Polya_svm (Cheng, et al., 2006)** | Support vector machine (SVM) | pA | 1 | Human | 100, +100 | 1) 29283 human genomic sequences surrounding the poly(A) sites (300 to +300 nt) in the polyA_DB database.<br>2) Negative sequences generated by a first-order Markov chain model derived from the positive sequences. | Position-specific scoring matrices (PSSMs) to score 15 cis regulatory elements. | Competing tools: polyadq, LDA, QDA<br>Indicators: SN, SP, CC<br>Results:<br>Better than polyadq, linear discriminant analysis (LDA), and quadratic discriminant analysis (QDA) on the test data.<br>SN=37.2-71.0%, SP=74.6-96.7%, CC=0.364-0.572. | Perl | https://exon.apps.wistar.org/polya_svm/ |
| 12 | **PolyApred (Ahmed, et al., 2009)** | SVM | NNUANA-like 13 variants | 1 | Human | -100, +100 | 1) A positive dataset containing 2327 sequences, each sequence 206 nt long having a poly(A) signal at the center (101 to 106 nt). A negative dataset containing 2333 sequences, each sequence 206 nt long extracted from coding regions having AATAAA at the center (101 to 106 nt).<br>2) **[ERPIN dataset]** and **[PSM dataset]**.<br>3) 50 sequences for each variant of poly(A) signal from Human ATD release 2. | Different types of nucleotide frequencies and binary pattern, split nucleotide frequency, 2$^{nd}$ order of dinucleotide frequency. | Competing tools: ERPIN, Polyadq, Polya_svm, Poly(A) Signal Miner<br>Indicators: SN, SP, MCC<br>Results: SN=57.0%, SP=75.8-95.7%, MCC=0.72. | Web server | http://www.imtech.res.in/raghava/polyapred/ |
| 13 | **POLYAR (Akhtar, et al., 2010)** | Linear discriminant model | pA | 1 | Human | -300, +300 | **[POLYAR dataset]:**<br>1) polya_DB and GenBank annotation of human genome (Build 34.2), with 29281 sequences of 600 bp length.<br>2) 28761 sequences were divided into 20225 PAS-strong, 6475 PAS-weak and 2061 PAS-less pAs.<br>3) Negative: 21,350 non-redundant sequences ("H2H" negative dataset); CDSs of 17600 human genes, 19600 | PAS-motif, [-40: -1], CS-motif, GU/U-motif, [+1: +50], upstream pentamer composition, downstream pentamer composition, PAS-CS distance, CS-GT distance. | Competing tools: polydq<br>Indicators: SN, SP<br>Results: For PAS-strong, compared with polya_svm (SN=65.7%, SP=51.7%), polydq (SN=53.7%,sp=70.4%), POLYAR has SN=80.8%, SP=66.4%. SN =13.5-80.8%, SP=14.766.4% across PAS-strong/-weak/- | Web server | - |

| | | | | | | | | | | | |
|---|---|---|---|---|---|---|---|---|---|---|---|
| | | | | | | | human intronic sequences and 3748 5' UTR regions, and generated 8261 randomized pA regions. | | less. | | |
| 14 | **(Chang, et al., 2011)** | SVM | 13 variants | 1 | Human | -125, +125 | 1) 33,745 positive sequences from PolyA_DB 2.<br>2) [ **ERPIN dataset]**<br>3) Negative: 6,000 sequence included randomized poly(A) regions (produced by randomizing the sequence surrounding a pA), 313,454 human mRNA coding sequences (CDS), 25,700 human 5' UTR and randomized genome sequences. | K-mer nucleotide patterns (k = 1, 2, 3), possible RNA secondary structures identified by Sfold, RNAfold, RNAMotif/ | Competing tools: Polya_svm<br>Indicators: SN, SP, CC<br>Results: Polya_svm: SN=54.92% (positive datasets); SP=80.12-93.84%, CC=0.33-0.394 (negative datasets), Chang et al.: SN=56.12% (positive datasets); SP=75.27%-94.28%, CC=0.288-0.403 (negative datasets). | - | - |
| 15 | **Dragon PolyA Spotter (Kalkatawi, et al., 2012)** | Artificial neural network (ANN), random forest (RF) | 12 variants | 1 | Human | -100, +100 | **[DPS dataset]**<br>1) 14799 positive sequences for 12 motif variants from human mRNA sequences database (http://hgdownload.cse.ucsc.edu/goldenPath/ hg19/bigZips/).<br>2) Negative records were selected from human chromosome 21 randomly (Beaudoing et al., 2000).<br>All sequences were deposited in http://cbrc.kaust.edu.sa/dps/ code/DataToBuildModel.tar.gz. | Thermodynamic, physico-chemical and statistical characteristics. | Competing tools: polyadq, POLYAR, poly_svm<br>Indicators: SN, SP, ACC<br>Results: ANN-average (overall) SN=89.62% (SD=4.31%, 80.55%-95.18%), SP=87.81% (SD=3.35%,83.57-93.37%), ACC=88.72%(SD=3.43%,82.06-94.27%); RF-average SN=86.67% (SD=2.65%, 83.1-93.2%), SP=93% (SD=1.40%, 91.1-95.6%), ACC= 89.84%(SD=1.75%, 88-94.4%). | C++, weka | http://cbrc.kaust.edu.sa/dps |
| 16 | **(Xie, et al., 2013)** | HMM and SVM | 12 variants | 1 | Human | -100, +100 | 1) **[DPS dataset]** | Using HMMs as a probabilistic generative model for DNA sequences, and developing an efficient spectral algorithm for extracting latent variable information from these models. | Competing tools: -<br>Indicators: FNR, FPR, error rate<br>Results: HMM average error rate=14.42%, false-negative rate=16.26%, false-positive rate =12.59%;<br>RF average error rate=19.19%, false-negative rate=18.83%, false-positive rate =19.48%. | Python | https://sfb.kaust.edu.sa/Pages/Software.aspx |
| 17 | **Omni-PolyA (Magana-Mora, et al., 2017)** | Combined classifier | 12 variants | 1 | Human | -100, +100 | 1) **[DPS dataset]**<br>2) The GENCODE PolyA feature annotation Release 19 (GRCh37.p13).<br>3) **[Omni dataset]:** 18,786 sequences with true PAS were extracted for the 12 most frequent PAS variants in human. For each PAS variant, the same number of pseudo-PAS sequences was generated from human chromosome 21 after excluding all the true PAS sequences contained in that chromosome. | Omni-PolyA feature set with 218 numeric values: mono-nucleotide and di-nucleotides frequencies in particular regions of the genomic DNA sequences, i.e., downstream, upstream, and in-frame codons with respect to the PAS hexamer, among others; novel and more specific features making use of the entropy and positional information gain of the nucleotide content to determine the most relevant sequence positions; a sequence score derived from 2-mer weight matrices. | Competing tools: Dragon PolyA Spotter, Xie et al.<br>Indicators: FPR, FNR, error rate<br>Results: For the DPS dataset(Kalkatawi, et al., 2012), average error rate= 12.99% (5.85-16.23%), FPR=12.05% (9.25-13.60%), FNR=13.93% (4.72-17.91%) across 12 variant (~24.5% error rate for PAS-strong variants);<br>average error rate=12.50% (5.45-14.49%), FPR=11.08% (6.67-15.14%), FNR=14.11% (4.23-18.84%) across 12 variant after pooling the PAS-weak variants sequences to expand the training data. | MATLAB | - |

**PAS/pA**: whether the tool predicts poly(A) site (pA) or poly(A) signal(s) (PAS) | **Site#**: how many pAs or PASs are predicted by the tool ("Multi" means multiple).

**Species**: species for the model training (please note that this is based on the data used in the respective study, while the tool may be trained or used for other species).

**Range**: the range of the sequence surrounding a pA or PAS used for model training or prediction (e.g., "-100, +200" means that the sequence upstream 100 nt to downstream 200 nt surrounding a pA or PAS is used; "-" means the sequence length is not fixed in the tool). |

**Data**: data used in the respective study for model training and testing | **Features**: features used in the model | **Performance**: summary of performance described in the respective study.

**Code**: programming language used in the tool | **Availability**: URL of the tool if available or "-" if no URL or the URL provided in the respective study is not accessible.

**Abbreviation:** pA, poly(A) site; PAS, poly(A) signal; SN, sensitivity; SP, specificity; FNR, false negative rate; FPR, false positive rate; ACC, accuracy; MCC, Matthews correlation coefficient; AUC, area under curve.

**Note** [1] If a dataset is used by multiple studies, we name the dataset after the study in which the dataset first appeared, e.g., '**[ERPIN dataset]**' means that the dataset was constructed in the study of ERPIN and was also used by other studies.
