## Supplementary material for "A Survey on Methods for Predicting Polyadenylation Sites from DNA Sequences, Bulk RNA-seq, and Single-cell RNA-seq": File S2

### File S2. Methods for predicting poly(A) sites or poly(A) signals from DNA sequences based on deep learning models.

| | Method | Model | PAS/pA | Site# | Species | Sequence length | Data | Features | Performance | Code | Availability | Additional functions |
|---|---|---|---|---|---|---|---|---|---|---|---|---|
| 1 | **deepPolyA (Gao, et al., 2018)** | Convolution neural network (CNN) | pA | 1 | Arabidopsis | -131, +31 | 1) **[PASS dataset]**.<br>2) Negative samples are generated by randomly sampling sequences from the Arabidopsis Information Resources (TAIR). | Encoded 4 × 162 bit matrix with regulatory variants, k-mer and structural elements. | Competing tools: SVM, Bayesian networks and Random Forest, DanQ, DeepSEA and VGG<br>Indicators: ACC, Recall, SP, MCC, AUC, F-score<br>Results: ACC=91.28%, Recall=93.01%, SP=89.51%, MCC=82.60%, AUC=97.06%, F score=91.51%. | Python | https://github.com/stella-gao/DeepPolyA | - |
| 2 | **Conv-Net (Leung, et al., 2018)** | CNN | pA strength | - | Human | -100, +100 | **[Conv-Net dataset]:**<br>The final pA atlas contains 19320 3' UTR regions with two or more pAs from genes in the hg19 assembly for a total of 92 218 sites in seven distinct human tissues, including the brain, breast, embryonic stem cells, ovary, skeletal muscle, testis and two samples of naïve B cells. | Cis-elements, RNA binding motifs, all possible 1-4 nt k-mers counts and nucleosome positioning features. | Competing tools: -<br>Indicators: AUC<br>Results: AUC=0.907 ± 0.004 | Python | - | Analyzing relative dominance of pAs in human tissues; predicting the consequence of nucleotide substitutions on PAS strength; assessing the effects of anti-sense oligonucleotides to alter transcript abundance. |
| 3 | **DeeReCT-PolyA (Xia, et al., 2019)** | CNN | 12 variants | 1 | Human, mouse | -100, +100 | 1) **[DPS dataset]**<br>2) **[Omni dataset]**<br>3) **[DeeReCT-PolyA dataset]**: 46224 genomic sequences from C57BL/6J (BL) and 40230 sequences from SPRET/EiJ (SP) mouse strains, respectively. | Cis-elements and variants | Competing tools: RF, HSVM, Omni-polyA.<br>Indicators: error rate<br>Results: For DPS dataset: Error Rate=11.81% for AATAAA; 4.59-11.8% across 12 variants (average=9.57%).<br>For Omni dataset: Error rate = 21.99% (17.82-27.76 across 12 variants, average=22.64%). | Python | https://github.com/likesum/DeeReCT-PolyA | - |
| 4 | **DeepPASTA (Arefeen, et al., 2019)** | CNN and recurrent neural network (RNN) | pA | 1 | Human | -100, +100 | 1) Postive data from PolyA-Seq data in (Derti et al., 2012);<br>2) **[Conv-Net dataset]** (Leung et al., 2018).<br>3) **[PLYAR dataset]** Akhtar et al., 2010). | 200 bp sequence and matched RNA secondary predicted by RNAshapes. | Competing tools: DeepPASTA, PolyAR, Dragon PolyA Spotter, DeepPolyA<br>Indicators: AUC, AUPRC<br>Results: AUC=0.972-0.958, AUPRC=0.875-0.962 | Python | https://github.com/arefeen/DeepPASTA | Tissue-specific pA prediction, including tissue-specific, tissue-specific relatively dominant pA and tissue-specific absolutely dominant pA. |
| 5 | **DeepGSR (Kalkatawi, et al., 2019)** | CNN | 16 variants | 1 | Human, mouse, bovine and fruit fly | -300, +300 | **[DeepGSR dataset]**:<br>20933, 18693, 12082, and 27203 true PAS data in total for the 16 PAS motifs | Sequence | Competing tools: -<br>Indicators: ACC<br>Results: A maximum ACC of 86.94% (representing a reduction of the relative error rate of 28.51%). | Python | https://doi.org/10.5281/zenodo.1117159 | Recognition of different genomic signals and regions (GSRs) in DNA, such as translation initiation sites (TIS). |
| 6 | **HybPAS (Albalawi, et al., 2019)** | Deep neural networks (DNNs) and | 12 variants | 1 | Human | -300, +300 | 1) Positive dataset: 37516 presumed true functional PAS sequences from GENCODE annotation for poly(A). | Three sets of features (signal processing-based, | Competing tools: Omni-PolyA,<br>Indicators: ACC<br>Results: HybPAS outperforms | Python, Matlab | https://github.com/EMANG-KAUST/PolyA_Prediction_LRM_DNN | - |

| | | | | | | | | | | | | |
|---|---|---|---|---|---|---|---|---|---|---|---|---|
| | | logistic regression models (LRMs) | | | | | 2) Negative dataset: 37 516 pseudo-PAS sequences sampled from the genome. | position weight matrix (PWM)-based, and statistics-based). | Omni-PolyA, reducing the classification error for different PAS hexamers by up to 57.35% for 10 out of 12 PAS types, with Omni-PolyA models being better for two PAS types. For the most frequent PAS types, 'AATAAA' and 'ATTAAA', HybPAS reduced the error rate by 35.14% and 34.48%, respectively. On average, HybPAS reduces the error by 30.29%. | | | |
| 7 | **APARENT (Bogard, et al., 2019)** | CNN | motifs | 1 | Human | -90, +90 | 1) Random sample of 120,000 sequences from the Alien1 library. | 1-Hot-Coded matrix | - | Python | https://github.com/johli/aparent | Predicting APA, visualizing motif interactions, predicting cleavage distribution, predicting APA across different tissue and cell types, predicting SNVs linked to APA misregulation. |
| 8 | **SANPolyA (Yu and Dai, 2020)** | DNN with attention mechanism | 18 variants | 1 | Human, mouse | -100, +100 | 1) **[DPS dataset]**<br>2) **[Omni dataset]**<br>3) **[DeepGSR dataset]**<br>4) **[DeeReCT-PolyA dataset]** | Four DNA sequences representation methods, including mononucleotide, dinucleotides, trinucleotides and embedding. | <u>Competing tools:</u> DPA, HMM-SVM, Omni-PolyA, DeeReCT-PolyA<br><u>Indicators:</u> error rate<br><u>Results:</u> DPS dataset: error rate 4.15 to 8.69 for different PAS motifs; omni-human dataset: 10.88 to 19.00; mouse datasets: 12.85 to 16.92. | Python | https://github.com/yuht4/SANPolyA | - |
| 9 | **Poly(A)-DG (Zheng, et al., 2020)** | CNN-Multilayer Perceptron network | 13 variants | 1 | Human, mouse, bovine, rat | -300, +300 | 1) **[Omni dataset]**<br>2) **[DeepGSR dataset]**<br>3) **[DeeReCT-PolyA dataset]**<br>4) Rat Poly(A) dataset constructed by the authors. | Both motif features and general sequence information. | <u>Competing tools:</u> DeeReCT-PolyA, DeepPolyA, SanPolyA.<br><u>Indicators:</u><br><u>Results:</u> | Python | https://github.com/Szym29/PolyADG | Visualization of the motif features that the convolution filters most likely to capture; predicting the number of PAS on whole genome. |
| 10 | **PASNet (Guo, et al., 2021)** | Self-attentive gated convolutional highway networks | 16 variants | 1 | Human, mouse, bovine and fruit fly | -300, +300 | 1) For the 16 different PAS motifs of the Human, Mouse, Bovine, and Fruitfly, the number of 16 polyadenylation motifs for PAS is 20933, 18693, 12082, and 27203 biological sequences respectively.<br>2) **[DeepGSR dataset]** | Split each biological sequence into a k-mer sequence corpus by using the window of length k (k = 5) | <u>Competing tools:</u> DeepPASTA, DeepPolyA, DeepGSR, DPS, HMM-SVM and Omni-PolyA<br><u>Indicators:</u> error rate, SN, SP, ACC<br><u>Results:</u> PASNet achieves the improvement with the error rate decreasing by 4% and 1.02% respectively on average.<br>For the four species, SN=82.46% to 88.22%, SP=83.87% to 87.93, ACC=83.51% to 87.63%. | Python | https://gitlab.com/guoyb/pasnet | - |

**Additional functions**: functions in addition to pA/PAS prediction provided by the tool.
Other columns are the same as in File S1.
