## Supplementary material for "A Survey on Methods for Predicting Polyadenylation Sites from DNA Sequences, Bulk RNA-seq, and Single-cell RNA-seq": File S3

### File S3. Methods for predicting poly(A) sites from RNA-seq.

| | Method | Model | pA # | Species | Smp# | Data | Performance | Input | pA Anno | Annotation | Code | Availability | Additional functions |
|---|---|---|---|---|---|---|---|---|---|---|---|---|---|
| ***Methods that interrogate non-templated poly(A)-capped reads*** | | | | | | | | | | | | | |
| 1 | **KLEAT (Birol, et al., 2015)** | Alignment and poly(A) tails | ≥1 | Human | 1 | **[RNA-PET dataset]**<br>Matched ENCODE RNA-seq and RNA-PET libraries of three cell lines (H1-hESC, A549 and MCF-7). | Competing tools: -<br>Indicators: PPV, TP<br>Results: KLEAT has over 90% positive predictive value when there are at least three RNA-seq reads supporting a poly(A) tail and requiring at least three RNA PET reads mapping within 100 nucleotides as validation.<br>TP: 75%~93% | An alignment file in BAM format;<br>Annotation and EST database in GTF format (optional). | N | N | Python | https://www.bcgsc.ca/resources/software/kleat | - |
| 2 | **ContextMap 2 (Bonfert and Friedel, 2017)** | Context-based mapping approach | ≥1 | All | 1 | 1) **[RNA-PET dataset]**<br>2) SAPAS data on MCF-7 from (Fu, et al., 2011).<br>3) RNA-seq data from (Pickrell, et al., 2010);<br>4) HSV-1 sequencing data. | Competing tools: KLEAT<br>Indicators: SN, PPV<br>Results: SN=0.188~0.381; PPV=0.826~0.96 (ENCODE)<br>SN=0.143~0.153; PPV=0.802~0.812 (SAPAS data)<br>SN=0.73-0.91; PPV=>0.84 (HSV-1)<br>SN=0.813; PPV=0.78 (Pickrell data) | Reference genome in FASTA format;<br>RNA-seq reads in FASTA or FASTQ format;<br>Reference annotation in GTF format (optional). | N | N | Java | https://www.bio.ifi.lmu.de/software/contextmap | - |
| ***Methods based on transcript assembly*** | | | | | | | | | | | | | |
| 3 | **PASA (Campbell, et al., 2006)** | An all-vs-all comparison within assembled transcript clusters | ≥1 | All | 1 | All publicly available ESTs and mRNA sequences for Oryza sativa and Arabidopsis thaliana from GenBank and DDBJ databases. | - | Reference genome in FASTA format;<br>ESTs or de novo RNA-seq assemblies in FASTA format;<br>Reference annotation in GFF3 format (optional). | N | N (optional) | Perl, C++ | https://github.com/PASApipeline/PASApipeline | Identify gene structures, antisense transcript, splicing variation. |
| 4 | **3USS (Le Pera, et al., 2015)** | Transcript assembly by Cufflinks | ≥1 | Mouse | 1 or 2 | - | Competing tools: -<br>Indicators: -<br>Results:The tool reports as putative novel the 3' UTRs not annotated in available databases. | Reference genome in FASTA format;<br>Genome annotation in GTF format;<br>One or two RNA-seq transcriptome assembly files. | N | Y | Web-server | https://www.biocomputing.it/3uss_server (not accessible) | - |
| 5 | **ExUTR (Huang and Teeling, 2017)** | Transcript assembly | - | Mammals | 1 | RNA-seq data: Bat Blood SRX763357; Cow Brain SRX764721; Mouse Kidney SRX1603138; Pig Blood SRX242932; Rat Brain SRX471401; Human Liver ERX1217498; Human Liver ERX1217498; Arctic fox Mixed ERX632794; Spiny mouse Brain SRX1818436; Long-haired mouse Kidney SRX663111; Grey wolf Blood SRX1713277; Sika deer Antler ERX024230 ERX028792. | Competing tools: GETUTR, 3USS<br>Indicators: TP<br>Results: TP: reference-based=89.0%~94.8%; *de novo*=79.5%~86.1%. | RNA-seq assemblies in FASTA format;<br>ORF sequences in FASTA format;<br>3' UTR database in FASTA format. | Y | N | Perl | https://github.com/huangzixia/ExUTR | Transcriptome assembly, ORF prediction and 3' UTR retrieval. |
| 6 | **Scripture (Guttman, et al., 2010)** | Transcript assembly, a statistical segmentation approach | - | All | 1 | RNA-Seq data: mouse embryonic stem cells (ESC), mouse neural progenitor cells (NPC) and mouse lung fibroblasts (MLF) (GSE20851). | Scripture accurately reconstruct most expressed, annotated protein coding genes, at a broad range of expression levels, as well as uncover many new isoforms in the protein coding transcriptome. | An alignment file in SAM format;<br>Reference genome in FASTA format;<br>A chromosome size file | N | N (optional) | Java | https://software.broadinstitute.org/software/scripture/ | Build a transcriptome ab initio. |
| ***Methods that rely on prior annotations of pAs*** | | | | | | | | | | | | | |
| 7 | **QAPA (Ha, et al.,** | Build 3′ UTR reference | ≥1 | Human, mouse | 1 | 1) Synthetic RNA-seq dataset with two conditions. | Competing tools: GETUTR, Roar, DaPars<br>Indicators: CC, AUC | Gene annotation from Biomart or GENCODE; | Y | Y | Python, R | https://github.com/morrislab/qapa | Calculation of relative usage of |

| | | | | | | | | | | | | | |
|---|---|---|---|---|---|---|---|---|---|---|---|---|---|
| | **2018)** | library by using GENCODE and incorporating pA annotations | | | | 2) **[Liu HEK293 dataset]**: RNA-seq and 3′-seq data from HNRNPC knockdown and METTL3 knockdown in HEK293T cells (Gruber et al. 2016; Liu et al. 2015).<br>3) Human brain and skeletal muscle from RNA-seq (Parsons et al. 2015) and 3'seq (Lianoglou et al. 2013). | Results: PPAU's CC=0.70 (HEK293, RNAseq~A-seq2)<br>[PPAU, Proximal Poly(A) Usage]<br>QAPA AUC = 0.88; the AUC for Roar, DaPars, and GETUTR are 0.66, 0.65, and 0.62, respectively. | Poly(A) site annotation (optional);<br>Reference genome (fasta);<br>Output from Sailfish or Salmon for RNA-seq. | | | | | alternative 3' UTR isoforms based on transcript-level abundance. |
| 8 | **PAQR (Gruber, et al., 2018)** | Obtained poly(A) site from PolyAsite database (priori annotations) | ≥1 | Human | 2 | 1) RNA-seq of CFIm 25 knock-down in HeLa cells ( Masamha et al. 2014).<br>2) **[Liu HEK293 dataset]**<br>3) **[Gueroussov HEK293 dataset]**: PTBP1/2 knock-down in HEK293 cells (Gueroussov et al. 2015) | Competing tools: DaPars<br>Indicators: -<br>Results: Compared with DaPars, PAQR-based quantification of pA use led to much more reproducible HNRNPC binding motif activity and more significant difference of mean z-scores between conditions (−22.92 with PAQR-based). | Alignment files in BAM format;<br>Known prior poly(A) sites in bed format | Y | Y | Python, R | https://github.com/zavolanlab/PAQR2 | - |
| 9 | **APA-scan (Fahmi, et al., 2020)** | Read coverage estimation and APA identification. | ≥1 | Human mouse | 2 | **[MEF dataset]** | Competing tools: -<br>Indicators: -<br>Results: qPCR experiments for Srsf3 and Rpl22 transcripts. | Refseq annotation,<br>Reference genome (fasta);<br>alignment files of RNA-seq in BAM. | Y | Y | Python | https://github.com/compbiolabucf/APA-Scan | Visualization of 3' UTR APA. |
| ***Methods that infer pAs by detecting significant changes in RNA-seq read density*** | | | | | | | | | | | | | |
| 10 | **GETUTR (Kim, et al., 2015)** | Read density fluctuations , kernel density estimation | ≥1 | All | 1 | 1) RNA-seq and 3P-seq of four wild-type cell lines, HeLa, HEK293, Huh7,and IMR90 (Nam et al. 2014). | Competing tools: -<br>Indicators: -<br>Results: Predicts >30,000 sites from 11,902~13,079 genes using RNA-seq data, among which 8597~12,151 (26.0~38.6%) validated by 3P-seq. | An alignment file of RNA-seq in BAM or BED format;<br>Reference annotation in GenePred or GTF format. | N | Y | Python | http://big.hanyang.ac.kr/GETUTR/manual.htm | - |
| 11 | **EBChangePoint (Zhang and Wei, 2016)** | Empirical Bayes change-point model | 1 | Human | 2 | 1) Simulated data.<br>2) Real RNA-seq data of melanoma cell migration (Flockhart et al. 2012). | Competing tools: -<br>Indicators: FDR | 3'UTR and junctions in BED;<br>alignment files of RNA-seq in BAM. | N | Y | Java | http://ebchangepoint.sourceforge.net/ | Identifying 30/50 AS events |
| 12 | **IsoSCM (Shenker, et al., 2015)** | Read density fluctuations (change point), a Bayesian model | ≥1 | All | 1 | 1) Simulated data.<br>2) **[PolyA+ mouse dataset]**: Real RNA-seq data: nine mouse tissues analyzed in triplicates (GSE41637) (Merkin et al. 2012). | Competing tools: Cufflinks, Scripture<br>Indicators: -<br>Results: IsoSCM made 4785 annotations, compared with 2450 by Cufflinks and 1827 by Scripture.<br>IsoSCM identified 2718 novel termini supported, compared with 1914 identified by Cufflinks and 941 identified by Scripture. | Coverage from bulk RNA-seq;<br>GTF genome annotation. | N | N | Java | https://github.com/shenkers/isoscm | Detect APA site switching. |
| 13 | **DaPars (Xia, et al., 2014); (Updated version: DaPars2 (Feng, et al., 2018; Li, et al., 2021))** | Read density fluctuations, a regression model | 1 | Human | 2 | 1) Simulated 1,000 genes in tumor and normal.<br>2) **[MAQC dataset]**: RNA-seq data and PolyA-seq data based on the same Human Brain Reference and the Universal Human Reference (UHR) MAQC samples (Bullard et al. 2010; Derti et al. 2012; Consortium et al. 2006). | Competing tools: Cufflinks<br>Indicators: AUC<br>Results: AUC =0.762~ 0.985 (simulated data). From the comparison between brain and UHR, 60% of 372 DaPars predicted APA events could be strongly supported by PolyA-seq. Both PolyA-seq and DaPars reported longer 3' UTRs in brain than in UHR in >94% dynamic APA events. | Coverage from bulk RNA-seq;<br>GTF genome annotation. | N | Y | Python | https://github.com/ZhengXia/DaPars | Detect APA site switching. |
| 14 | **APAtrap (Ye, et al., 2018)** | Read density fluctuations, a mean squared error model | ≥1 | Human, Arabidopsis | 2 | 1) Simulated 1,000 genes with 2 isoforms and 1-4 isoforms.<br>2) **[MAQC dataset]**.<br>3) Arabidopsis thaliana using RNA-seq data from control and mild drought conditions. | Competing tools: DaPars, ChangePoint<br>Indicators: AUC<br>Results: AUC: APAtrap=0.95, DaPars=0.90, ChangePoint=0.67 (simulated 1 isoform); APAtrap=0.94, DaPars=0.78 (simulated 1-4 isoforms). | Coverage from bulk RNA-seq;<br>GTF genome annotation. | N | Y | R | https://sourceforge.net/projects/apatrap/ | Detect APA site switching. |
| 15 | **TAPAS** | Read density | ≥1 | Human | 2 | 1) Simulated data. | Competing tools: Cufflinks, IsoSCM, GETUTR, | Coverage from bulk RNA-seq; | N | Y | R | https://github.com | Detect APA site |

| | | | | | | | | | | | | | |
|---|---|---|---|---|---|---|---|---|---|---|---|---|---|
| | **(Arefeen, et al., 2018)** | fluctuations, PrunedExactLinearTime method (Killick, et al., 2012) | | , mouse | | 2) **[PolyA+ mouse dataset]** mouse brain.<br>3) **[MAQC dataset]**<br>4) PAS-Seq data from NCBI (GSE25450) of mouse ES (embryonic stem) and NPS (Neuropeptide S) and neuron cells (Shepard et al. 2011). | Indicators: SN, Precision<br>Results: SN:<br>TAPAS=77.61%~86.84%; Cufflinks=69.56%~77.21%; IsoSCM=54.25%~67.15%; GETUTR=71.30%~71.30%;<br>Precision (simulated data):<br>TAPAS=86.70%~88.45%; Cufflinks=58.25%~65.80%; IsoSCM=41.88%~49.30%; GETUTR=30.49%~31.3%;<br>Precision (mouse brain):<br>TAPAS=30.84%~36.15%, Cufflinks=30.84%~11.13%, IsoSCM=17.51%~21.17%, GETUTR=4.95%~11.10%;<br>Precision (mouse neuron): TAPAS=77.88%~86.84%, Cufflinks=17.26%~24.18%, IsoSCM=47.38%~54.89%, GETUTR=9.95%~24.57% | GTF genome annotation. | | | | /arefeen/TAPAS | switching. |
| 16 | **mountainClimber (Cass and Xiao, 2019)** | Calling change points in each transcript unit of each sample | ≥1 | Human | 1 | 1) Simulation data with Flux Simulator.<br>2) **[MAQC dataset]**. | Competing tools: IsoSCM, MISO<br>Indicators: Precision<br>Results:<br>1) For simulation data, mountainClimber outperformed IsoSCM in terms of recall, while the two methods yielded similar precision. mountainClimber outperformed the other two methods (MISO, IsoSCM) in terms of recall, especially for TandomAPA sites.<br>2) For real data, mountainClimber achieved higher precision than IsoSCM regardless of the window size, w, around PolyA-seq sites used to define true positives.<br>Similarly, mountainClimber outperformed IsoSCM in terms of TSS precision and predicted more change points with high fold changes near FANTOM CAT-predicted 5'ends. | Alignment BAM file,<br>chromosome sizes,<br>GTF genome annotation. | N | Y | Python | https://github.com/gxiaolab/mountainClimber | Recognizes multiple transcription start sites or APA sites in a transcript. |
| 17 | **IPAFinder (Zhao, et al., 2021)** | Read density fluctuations, considering junction-spanning reads | ≥1 | Human and mouse | 2 | 1) TCGA RNA-seq BAM files for tumor and matched normal samples obtained from the Genomic Data Commons (GDC) Data Portal.<br>2) Data of 3′-seq and RNA-seq: normal immune cells and malignant B cells from patients with chronic lymphocytic leukemia (CLL) (Singh et al. 2018; Lee et al. 2018).<br>3) **[Gueroussov HEK293 dataset]**<br>4) **[Liu HEK293 dataset]**<br>5) YTHDC1 knockdown and SRSF3 knockdown in HeLa cells (Xiao et al. 2016).<br>6) U2AF1 knockdown in HFF cells (Yao et al. 2020).<br>7) U1 inhibition in HeLa cells (So et al. 2019).<br>8) Synthetic RNA-seq. | Competing tools:-<br>Indicators: -<br>Results:<br>1) Use IPAFinder to infer and quantify IPA sites based on the synthetic RNA-seq data set.<br>2) Compare results of IPAFinder with 330 recurrent CLL-IpA events were obtained from the data sets of Lee et al. 2018. | Bam from bulk RNA-seq;<br>GTF annotation. | N | Y | Python | https://github.com/ZhaozzReal/IPAFinder | Detect dynamic changes of IPAs |
| ***Methods based on machine learning models*** | | | | | | | | | | | | | |
| 18 | **IntMAP (Chang, et al., 2018)** | A constrained probabilistic model by combining | ≥1 | Mouse | 1 | **[MEF dataset]**:<br>Generated RNA-seq (SRP056624) and PAS-seq (PRJNA436720) from wild type and Tscl-/- MEF cells. | Competing tools:-<br>Indicators: -<br>Results: IntMAP found more (957) 3'UTR shortening compared with RNA-seq (846) and PAS-seq (843) method, respectively. | Processed RNA-seq (mat format) from the output by TopHat<br>Processed PAS-seq (mat format) from output by Bowtie and peak calling. | Y (PAS-seq) | Y | Matlab | http://compbio.cs.umn.edu/IntMAP/ | Detect APA site switching. |

| | | | | | | | | | | | | | |
|---|---|---|---|---|---|---|---|---|---|---|---|---|---|
| | | RNA-seq read alignments and PAS-seq peak callings | | | | | Among 975 genes, 592 and 370 genes overlapped with the PAS-Seq and RNA-Seq analyses, respectively.<br>Importantly, IntMAP identified 280 new 3'-UTR APA events, which were not reported in the union of two datasets and validated by qRT-PCR. | | | | | | |
| 19 | **TECtools (Gruber, et al., 2018)** | Constructing gene models | ≥1 | Human | 1 | **[Liu HEK293 dataset]** | Competing tools: StringTie, Cufflinks<br>Indicators: -<br>Results: The median lengths of TECtool-predicted terminal exons in the two HEK293 RNA-seq samples were 732 and 632 nt, respectively, longer than StringTie (380 and 412 nt, respectively) and Cufflinks (199 and 232 nt, respectively).<br>TECtool made more reproducible predictions from replicate samples (40% of the union of predicted exons were identified from both replicates, compared with 30% for StringTie and 18% for Cufflinks). | Chromosomes in fasta format; annotation in GTF format; genome coordinates of 3'end processing sites in BED format; spliced alignments of mRNA-seq reads to the corresponding genome in BAM format. | Y | Y | Python | https://github.com/zavolanlab/TECtool | Generate new isoforms and can be applied for scRNA-seq. |
| 20 | **Terminitor (Yang, et al., 2020)** | Assembling RNA-seq data using RNA-Bloom and deep neural network to predict sequence | ≥1 | Human and mouse | 1 | 1) Poly(A) annotations: PolyA_DB version 3.1, APADB v2, APASdb, and PolyASite.<br>2) **[MAQC dataset]** | Competing tools: KLEAT, DeeReCT-PolyA and DeepGSR<br>Indicators: precision<br>Results: Termin(A)ntor trained on human sequences consistently outperforms all Competing tools in both metrics, except for one data point. The four models built by DeeReCT-PolyA and DeepGSR all have a precision lower than 50%, while precisions of the Termin(A)ntor human model (at probability = 0.5 cut-off) are 55.59% and 57.72% for UHR and HBR samples, respectively. | Alignment BAM file;<br>FASTA file;<br>annotation file (Transcript annotation file; Ensembl annotation file). | N | Y | Python | https://github.com/bcgsc/TerminAntor | - |
| 21 | **Aptardi (Lusk, et al., 2021)** | Bidirectional long short term memory recurrent neural network (biLSTM) | ≥1 | Human and mouse | 1 | A total of five unique datasets: **[MAQC dataset]**, 2nd HBR, UHR, BNLx, and SHR. (HBR55: Accession: PRJNA510978, SRA runs: SRR8360036-37; 2nd HBR and UHR56: Accession: PRJNA362835, 2nd HBR SRA runs: SRR5236425-30, UHR SRA runs: SRR5236455-60; BNLx and SHR71: Accession: GSE166117, BNLx: GSM5061950-52, SHR: GSM5061947-49 | Competing tools: APARENT, TAPAS<br>Indicators: precision, recall, F-measure<br>Results: 1) Using poly(A) site from PolyA-seq as standard, the average precision is 0.74, recall is 0.39 and F-measure=0.51 based on HBR dataset across five cross-validation.<br>2) The positive predictive value of APARENT was lower than that for aptardi for all databases at all base distance cutoffs.<br>3) Although the number of transcripts with a false positive 3' terminus was higher in the aptardi modified transcriptome compared with TAPAS, the positive predictive value was higher for the aptardi modified transcriptome. | Bam from bulk RNA-seq;<br>DNA sequence;<br>GTF annotation. | N | Y | Python | https://github.com/luskry/aptardi | Construct new transcript annotations. |

**pA#**: number of predicted pAs for a gene | **Smp#**: number of the required sample(s) | **pA anno**: whether the pA annotation is required (Y) or not (N) | **Annotation**: whether the genome annotation (GTF file) is required (Y) or not (N).
**Input**: input data for the tool | **Additional functions**: functions in addition to pA prediction provided by the tool. | Other columns or abbreviations are the same as in File S1.
**Abbreviation:** PPV, positive predictive value.
