## Supplementary material for "A Survey on Methods for Predicting Polyadenylation Sites from DNA Sequences, Bulk RNA-seq, and Single-cell RNA-seq": File S4

### File S4. Methods for predicting poly(A) sites from single-cell RNA-seq.

| | Method | Model | Protocol | Region | Resolution | Data | pA | Species | Performance | Code | Availability |
|---|---|---|---|---|---|---|---|---|---|---|---|
| ***Methods based on peak calling*** | | | | | | | | | | | |
| 1 | **scAPA (Shulman and Elkon, 2019)** | Peak calling | 3' tag | Gene and extended 3'UTR | Peak | 1) Mouse Brain scRNA-seq dataset (SRP135960) (Zeisel et al. 2018; La Manno et al. 2018).<br>2) Mouse T cells (GSE106264) (Pace et al. 2018).<br>3) **[Mouse sperm dataset]**: Mouse Sperm cells (GSE104556) (Lukassen et al. 2018).<br>4) Lung tumor (E-MTAB-6149) (Lambrechts et al. 2018). | N | Mouse human | Competing tools: -<br>Indicators: SN, SP<br>Results: For peaks identified on a training set which included 158 unimodal and 42 bimodal cases, the above criterion showed SN=81% (34 out of 42 cases) with no false positive splits (SP of 100%). | R | https://github.com/ElkonLab/scAPA |
| 2 | **polyApipe** | PolyA reads & peak calling | 3' tag | Whole genome | Pos | 1) The 'pbmc8k' dataset (https://support.10xgenomics.com/single-cell-gene-expression/datasets/2.1.0/pbmc8k) | N | All | -<br>This tool is presented in a F1000Research poster (not peer reviewed). | Python, R | https://github.com/MonashBioinformaticsPlatform/polyApipe |
| 3 | **Sierra (Patrick, et al., 2020)** | Peak calling | 3' tag | Gene and extended 3'UTR | Peak | 1) **[PBMCs]**: two human PBMC datasets (a 7k cell and 4k cell) from 10x Genomics.<br>2) Total non-cardiomyocyte cells (total interstitial population [TIP]) from uninjured (sham) hearts or hearts at 3 or 7 days following myocardial infarction (MI) surgery (Farbehi et al. 2019).<br>3) Enriched cardiac fibroblast lineage cells (Pdgfra-GFP +) also from MI and sham mouse hearts (Farbehi et al. 2019).<br>4) **[Tabula Muris atlas]**: Twelve tissues from Tabula Muris (The Tabula Muris Consortium. 2018). | N | All | Competing tools: -<br>Indicators: -<br>Results: There is a strong correlation between gene expression from CellRanger and expression of peaks from Sierra.<br>Sierra can detect multiple types of alternative mRNA isoform usage with a significant number of these corroborated by an independent bulk RNA-seq experiment.<br>The detected DTU (differential transcript usage) genes by Sierra is reproducible between PBMC 4k and PBMC 7k dataset. | R | https://github.com/VCCRI/Sierra |
| 4 | **scAPAtrap (Wu, et al., 2021)** | PolyA reads & peak calling | 3' tag | Whole genome | Pos & peak | 1) Arabidopsis root cells (Ryu et al. 2019).<br>2) Mouse intestinal organoids cells from CEL-seq (Grun et al. 2015).<br>3) **[Mouse sperm dataset]** | N | All | Competing tools: Sierra<br>Indicators: accurary<br>Results: accurary =91%. | R | https://github.com/BMILAB/scAPAtrap |
| 5 | **SAPAS (Yang, et al., 2021)** | PolyA reads & peak calling | 3' tag | 3' UTR | Pos | 1) scRNA-seq datasets of mESCs by CEL-seq2 and SCRB-seq (Ziegenhain et al. 2017).<br>2) scRNA-seq dataset of HEK293 cell line by Microwell-seq method and a 3′-seq dataset of HEK293 cell line (Han et al. 2018).<br>3) scRNA-seq data of six GABAergic neurons (Paul et al. 2017). | N | Mouse human | Competing tools: DaPars and QAPA<br>Indicators: -<br>Results: Compared the performance of SAPAS on quantifying single-cell APA profiles with DaPars and QAPA, using a CEL-seq2 dataset of PBMCs. | R | https://github.com/YY-TMU/SAPAS |
| 6 | **SCAPE (Zhou, et al., 2022)** | Probabilistic mixture model | 3' tag | 3' UTR | Pos | 1) Simulated data.<br>2) Mouse bone marrow dataset generated by this study.<br>3) Mouse cell atlas, mouse iPSC and human GBM (Glioblastoma) datasets from GSE108097 (Han et al. 2018), GSE103221 (Guo et al. 2019) and PRJNA5795936 (Bhaduri et al. 2020).<br>4) **[Mouse HSC dataset]**: Mouse 3'-seq dataset of ex vivo isolated mouse multipotent steady state hematopoietic stem cells (sHSC) and proliferating HSCs (16h pIC; pHSC) downloaded from PRJEB29693 (Sommerkamp et al. 2020). | N | All | Competing tools: Sierra, scAPA, scAPAtrap, SCAPTURE and MAAPER<br>Indicators: -<br>Results: Identified 31 558 sites from 36 mouse organs, 43.8% (13 807) of which were novel. | Python | https://github.com/LuChenLab/SCAPE |
| 7 | **ReadZS (Meyer, et al., 2021)** | Nonparametric statistical approach | 3' tag | Whole genome | Peak | 1) 57 cell types in the lung and blood (Travaglini et al. 2020). | N | All | Competing tools: Sierra<br>Indicators: -<br>Results: Over 90% of genes detected by ReadZS were not called by Sierra. | Nextflow | https://github.com/salzmanlab/ReadZS |

| | | | | | | | | | | | |
|---|---|---|---|---|---|---|---|---|---|---|---|
| ***Methods that rely on prior annotations of pAs*** | | | | | | | | | | | |
| 8 | **SCAPTURE (Li, et al., 2021)** | Peak calling and deep learning | 3' tag | Whole genome | Pos | 1) Positive stringent PAS in three databse of poly_DB3/polyA-seq,/PolySite v2.0 or GENCODE (n=251,071); (9:1 as training and validataion); Negative: constructed with 200 bp sequences randomly extracted in intergenic regions without overlapping with any of the annotated PASs in three databases or GENCODE.<br>2) **[PBMCs]**: six PBMCs datasets.<br>3) Using priori pAs for training the deep learning model DeepPASS. | Y | Trained models on human and mouse | <u>Competing tools:</u> Sierra, scAPA, DeepPASTA and APARENT<br><u>Indicators:</u> AUC, SN, ACC, F1, SP<br><u>Results:</u> 1) Better than DeepPASTA and APARENT based on data1's validation data; AUC=0.991 for training dataset and AUC=0.988 for validation dataset. 2) Better than scAPA and Sierra with high sensitivity and ACC (data2) ACC=83.5%, SN=0.69,SP=0.66,F1=0.74 | Shell | https://github.com/YangLab/SCAPTURE |
| 9 | **MAPPER (Li, et al., 2021)** | Gaussian kernel and likelihood model | 3' tag | Each gene (3'UTR/intron/exon) | Pos | 1) Bulk RNA-seq: mouse; QuantSeq FWD and QuanSeq REV(as gold standard); 4-control/treated(AS/RC4/RC8);<br>2) scRNA-seq: two 10x genomics for placental samples, three stages (Vento-Tormo et al. 2018; Tsang et al. 2017).<br>3) Using priori pAs from PolyA_DB v3. | Y | Human, mouse, rat, and chicken as in PolyA-DB v3.0 | <u>Competing tools:</u> Sierra, scAPA<br><u>Indicators:</u> Precision, recall<br><u>Results:</u> 1) Based on bulk RNA-seq, Precision=90.7% (SD=2.0%); recall=75.5% (SD=2.5%); 2) Based on 10x genomic, MAAPER is more sensitive and accurate than Sierra and scAPA. | R | https://github.com/Vivianstats/MAAPER |
| 10 | **scUTRquant (Fansler, et al., 2021)** | Pipeline | 3' tag | Whole genome | Pos | 1) **[PBMCs]**: Three PBMCs datasets.<br>2) **[Mouse HSC dataset]**<br>3) Mouse ESC datasets (GSM3629847-8) (Guo et al. 2019).<br>4) **[Tabula Muris atlas]**<br>5) Mouse bone marrow (GSM2877127-32) (Dahlin et al. 2018).<br>6) Mouse brain (GSM3722100-115) (Ximerakis et al. 2019). | Y | Mouse human | <u>Competing tools:</u> -<br><u>Indicators:</u> CC<br><u>Results:</u>1) When comparing 3' UTR transcript counts obtained from bulk 3' end sequencing methods with scUTRquant, for FACS-sorted HSCs, Spearman's correlation=0.88; for ESC, Spearman's correlation = 0.72.<br>2) scRNA-seq derived 3′UTR isoform counts or isoform usage of biological replicates showed Spearman's correlation = 0.98 and correlation = 0.96 when performed by the same or by different laboratories, respectively | Snakemake | https://github.com/Mayrlab/atlas-mm |
| ***Other methods for predicting pAs from scRNA-seq*** | | | | | | | | | | | |
| 11 | **scDaPars (Gao, et al., 2021)** | Regression model for imputation; DaPars for pA identification. | 3' tag and full-length | 3' UTR | Pos | 1) 384 scRNA-seq libraries of individual human peripheral blood cells (PBMCs) sequenced by Smart-seq2 (Picelli et al. 2013) and a matched bulk RNA-seq library from a benchmark study by Ding et al. (Ding et al. 2020). | N | All | <u>Competing tools:</u> scAPA and Sierra<br><u>Indicators:</u> silhouette<br><u>Results:</u> 84% of poly(A) sites predicted from scRNA-seq data are within 100bp of those predicted in bulk, whereas only ~44% of randomly selected sites from 3' UTR regions are within 100bp of bulk predictions. ~66.2% of poly(A) sites predicted from scRNA-seq data also overlapped with annotated poly(A) sites complied from RefSeq, ENSEMBL, UCSC gene models and PolyA_DB. Compared with scAPA and Sierra, scDaPars showed higher silhouette coefficients. | R | https://github.com/YiPeng-Gao/scDaPars |
| 12 | **APA-seq (Levin, et al., 2020)** | Read1/2 mapping | CEL-seq | Gene and extended 3' UTR (downstream 5k bp of stop codon) | Peak | 1) C. elegans time-course data (GSE50548) sequenced using the CEL-Seq protocol and paired-end 100 bp sequencing mode. | N | C. elegans | <u>Competing tools:</u> -<br><u>Indicators:</u> -<br><u>Results:</u> Compared with a known repository of 3′ UTR annotations in C. elegans (41) and found highly concordant profiles, with 95% of sites corresponding to well-established annotated sites. | MATLAB, Python, Perl | https://github.com/yanailab/APA-Seq |

**Protocol**: protocol of the input scRNA-seq data | **Region**: the genomic region considered in pA prediction | **Resolution**: resolution of the detected pA, a position (pos) or a peak region (peak).
**Test data**: the data used in the study | **pA**: the tool is based on pA annotations (Y) or not (N).
Other columns are the same as in File S3.
