## Supplementary material for "A Survey on Methods for Predicting Polyadenylation Sites from DNA Sequences, Bulk RNA-seq, and Single-cell RNA-seq": File S5

### File S5. Tools or resources for APA analysis rather than pA prediction.

| Tool | Reference | Description |
|---|---|---|
| ***Methods for predicting tissue-specific pAs (task 2)*** | | |
| **polyA code** | (Weng, et al., 2016) | This tool predicts tissue-specific pAs based on a compendium of over 600 features. |
| **TSAPA** | (Ji, et al., 2018) | TSAPA predicts tissue-specific pAs in plants based on the machine learning model. |
| ***Methods for predicting dominant pAs (task 3)*** | | |
| **DeeReCT-APA** | (Li, et al., 2021) | An approach to predict APA usage based on bidirectional LSTM deep learning model, which outputs percentage scores representing the usage levels of all pAs of a gene. |
| ***Methods for predicting APA site switching (task 4)*** | | |
| **MISO** | (Katz, et al., 2010) | MISO was one of the first tools for differentially regulated AS/APA isoforms, using a probabilistic framework to quantify AS and APA. |
| **PHMM** | (Lu and Bushel, 2013) | PHMM is based on a two-state Poisson HMM to model the dynamic expression of 3′ UTRs, which considers a pA as transitions between two hidden states, e.g., the high and low expression states. |
| **ChangePoint** | (Wang, et al., 2014) | This is a change-point model using a likelihood ratio test for identifying switching of tandem pA usage based on the read coverage in RNA-seq data. |
| **roar** | (Grassi, et al., 2016) | roar quantifies and compares the ratio of short to long 3´ UTR isoforms between conditions, using annotated transcripts and pAs. |
| **SAAP-RS** | (Guvenek and Tian, 2018) | Significance Analysis of Alternative Polyadenylation using RNA-Seq (SAAP-RS) uses bulk RNA-seq, scRNA-seq and 3'READS to identify differential APA events, using an indicator called relative expression difference (RED) of the APA isoform to identify genes with significantly 3' UTR length changes between cell types. |
| **SCUREL** | (Burri and Zavolan, 2021) | SCUREL quantifies changes in 3'UTR length between groups of cells, including cells of the same type originating from tumor and control tissue. |
| **scDAPA** | (Ye, et al., 2019) | scDAPA employs a histogram-based approach and the Wilcoxon rank-sum test to measure the significance of the differential APA usage between conditions. |
| **satuRn** | (Gilis, et al., 2021) | satuRn is a fast and flexible quasibinomial generalized linear modelling framework for test differential usages of isoforms from the bulk RNA-seq, which is also suitable for scRNA-seq applications. |
| **scMAPA** | (Bai, et al., 2022) | scMAPA extends multiple modules of DaPars2 and combines a computational change-point algorithm and a statistical model to identify cell-type–specific APA genes from scRNA-seq, without assumptions on the read coverage shape. |
| ***Methods for other kinds of APA analysis (task 5)*** | | |
| **APAlyzer** | (Wang and Tian, 2020) | APAlyzer is a package for examining 3'UTR APA, intronic APA, and gene expression changes using RNA-seq data and annotated pAs in the PolyA_DB database. |
| **movAPA** | (Ye, et al., 2021) | movAPA is an R package that incorporates rich functions for preprocessing, annotation, and statistical analyses of pAs, identification of poly(A) signals, profiling of APA dynamics, and visualization. |
| **KAPAC** | (Gruber, et al., 2018) | KAPAC constructs 'impact maps' that show how motifs influence cleavage and polyadenylation. |
| **expressRNA's apa** | (Rot, et al., 2017) | expressRNA is a platform that provides exploratory tools for iCLIP, RNA-seq and 3´ seq data, including the software apa for the study of posttranscriptional processes. |
| **QuantifyPoly(A)** | (Ye, et al., 2021) | QuantifyPoly(A) can quantify genome-wide polyadenylation choices from 3' seq data, which reshapes 3' seq coverages into myriads of novel pA clusters. |
| **PolyA-miner** | (Yalamanchili, et al., 2020) | PolyA-miner creates a matrix of pA-sample from 3' seq data through non-negative matrix factorization for capturing gene expression patterns. |
| **acorde** | (Arzalluz-Luque, et al., 2022) | acorde is a pipeline based on percentile correlations that leverages bulk long reads and single-cell data to detect alternative isoform co-expression relationships, which can also detect and characterize genes with co-differential isoform usage across cell types. |
| ***Database resources of APA*** | | |
| **PolyA-Seq** | (Derti, et al., 2012) | PolyA-Seq contains filtered sites with normalized read counts, which is available via the UCSC Genome Browser (http://genome.ucsc.edu/). |
| **APASdb** | (You, et al., 2014) | APASdb catalogs pAs profiled by the sequencing alternative polyadenylation sites (SAPAS) method and details all the heterogeneous cleavage sites downstream of poly(A) signals. |
| **PolyA_DB 3** | (Wang, et al., 2018) | PolyA_DB 3 contains pAs in human, mouse, rat and chicken, which were mapped by the 3' region extraction and deep sequencing (3'READS) method. |
| **TC3A** | (Feng, et al., 2018) | The Cancer 3' UTR Atlas (TC3A) is a comprehensive resource of APA usage for 10,537 tumors across 32 cancer types. |
| **PolyAsite 2.0** | (Herrmann, et al., 2020) | PolyAsite 2.0 is an update of the PolyASite (https://polyasite.unibas.ch) resource of poly(A) sites, which was constructed from publicly available human, mouse and worm 3' end sequencing datasets. |
| **APAatlas** | (Hong, et al., 2020) | APAatlas includes APA events from 9475 samples across 53 human tissues Using the RNA-seq data from the Genotype-Tissue Expression project, and also provides information about traits and gene expression across tissues. |
| **PlantAPAdb** | (Zhu, et al., 2020) | PlantAPAdb catalogs pAs derived from 3' end data in six plant species. |
| **scAPAdb** | (Zhu, et al., 2021) | scAPAdb collects APA information from ~360 scRNA-seq experiments, covering six species including human, mouse and several other plant species. |
| **Animal-APAdb** | (Jin, et al., 2021) | Animal-APAdb provides APA information in 9244 samples of 18 animal species using DaPars2 and QAPA. |
| **3'aQTL-atlas** | (Cui, et al., 2021) | 3'aQTL-atlas provides 3' untranslated region (3'UTR) alternative polyadenylation (APA) quantitative trait loci (3'aQTLs), containing approximately 1.49 million SNPs associated with APA of target genes, based on 15,201 RNA-seq samples across 49 human Genotype-Tissue Expression (GTEx v8) tissues isolated from 838 individuals. |
| **TREND-DB** | (Marini, et al., 2020) | TREND-DB details the dynamic landscape of APA after depletion of ~170 proteins involved in various facets of transcriptional, co- and post-transcriptional gene regulation, epigenetic modifications. |
