## Supplementary material for "A Survey on Methods for Predicting Polyadenylation Sites from DNA Sequences, Bulk RNA-seq, and Single-cell RNA-seq": File S6

### Supplementary Text

### Predicting poly(A) sites (pAs) from bulk and single-cell RNA-seq

#### Human PBMCs data

We used bulk RNA-seq and scRNA-seq datasets of human peripheral blood mononuclear cells (PBMCs) for benchmarking [1]. The scRNA-seq PBMC data was sequenced by 10x Chromium (v2) and was downloaded from NCBI GEO (Accession nos. GSM3837156, GSM3837157, GSM3837158, GSM3837159, GSM3837170, GSM3837171, and GSM3837172). The matched bulk RNA-seq data was also downloaded from GEO (Accession no. GSM3839705). The 10x PBMC dataset contains 3362 cells. The cell type annotation was obtained from the original study, including nine cell types: CD4+T cell, CD8+ cytotoxic T cell, Nature killer cell, B cell, CD14+ monocyte, CD16+ monocyte, Dendritic Cell, and Plasmacytoid dendritic cell and Platelet.

#### Predicting poly(A) sites from bulk RNA-seq

We chose three representative tools for bulk RNA-seq, including DaPars2 [2], TAPAS [3], and Aptardi [4]. DaPars2 is the updated version of DaPars [5] that is probably the first and the most widely used tool for bulk RNA-seq. TAPAS generally obtained overall high sensitivity and accuracy according to previous benchmark studies [6, 7]. Aptardi is the latest tool based on deep learning model, using DNA sequences, RNA-seq and transcriptome assemblers.

Raw RNA-seq data were aligned to the hg38 reference genome using STAR

[8]. We followed the tutorial provided in the respective study for predicting pAs from bulk RNA-seq, using the BAM file generated by STAR and the genome annotation of hg38 as inputs.

#### Predicting poly(A) sites from scRNA-seq

We chose three representative tools for scRNA-seq, including Sierra [9], scAPAtrap [10] and SCAPTURE [11]. Sierra is one of the first tools for scRNA-seq based on peak calling, which is splice-aware and can identify pAs in introns. scAPAtrap can identify precise position of pAs by anchoring poly(A) reads and can predict pAs genome-wide, including intergenic pAs. SCAPTURE incorporates a deep learning model for pA filtering and uses annotated pAs from four databases for model training. Raw scRNA-seq data were aligned to the hg38 reference genome using Cell Ranger. We followed the tutorial provided in the respective study for predicting pAs from scRNA-seq, using the BAM file and the genome annotation of hg38 as inputs. For scAPAtrap, we retained peaks with at least 80 reads for downstream analyses.

#### Predicting poly(A) sites from DNA sequences using DeepPASTA

DeepPASTA [12] is one of the first tools using deep learning model for predicting pAs from DNA sequences. Because tools for DNA sequence are not applicable to RNA-seq data, here we extracted the upstream 100 nt and downstream 99 nt sequences of pAs (a total length of 200 nt) predicted by each of the RNA-seq tools, and then used DeepPASTA to classify each sequence as true (probability score>0.5) or false (probability score<=0.5). A sequence classified as true means that the middle position of the sequence is predicted as a real pA by DeepPASTA.

### Annotation of poly(A) sites

We used movAPA [13] for pA annotation and poly(A) signal analyses. Annotated

3′ UTRs were extended by 1000 bp to consider the effect of 3′ UTR extension. Each pA was annotated with its genomic region, such as 3′ UTR, CDS, intron, 5′ UTR and intergenic. The upstream 50 bp to downstream 25 bp region of each pA was scanned for poly(A) signal motifs including AATAAA and its 1-nt variants (TATAAA, CATAAA, GATAAA, ATTAAA, ACTAAA, AGTAAA, AAAAAA, AACAAA, AAGAAA, AATTAA, AATCAA, AATGAA, AATATA, AATACA, AATAGA, AATAAT, AATAAC, and AATAAG). Motifs within the upstream 50 bp to downstream 25 bp region of pAs predicted by each tool were identified by DREME [14] and the sequence logo of the most dominant motif was obtained.

### Collection of annotated poly(A) sites

Annotated poly(A) sites of human were downloaded from GENCODE v39 [15] (n=50,964, hg38), PolyA_DB 3 [16] (n=290,168, Ensembl v75, hg19) and PolyASite 2.0 [17] (n=569,005, Ensembl v96, hg38). Coordinates based on hg19 annotation were converted to hg38 using LiftOver (Genome: Ensembl v 102). Similar to [18], we then combined pAs from these databases, with the following priority: GENCODE v39 > PolyA_DB 3 > PolyASite 2.0. PAs within 24 bp of each other were grouped into pA clusters. Each cluster is considered as a non-redundant pA and the coordinate of the last pA in a cluster was considered as the coordinate of the cluster. Finally, a total of 676,424 non-redundant pAs were compiled for evaluation.
