## Supplementary material for "A Survey on Methods for Predicting Polyadenylation Sites from DNA Sequences, Bulk RNA-seq, and Single-cell RNA-seq": Table 1

**Table 1 Recommended tools for predicting pAs from DNA sequences, bulk RNA-seq, and single-cell RNA-seq**

*Note*: Tools were chosen based on criteria such as availability, function, ease of use, and popularity. pA, poly(A) site; ML, machine learning; SVM, support vector machine; GHMM, generalized hidden Markov model; CNN, convolution neural network; RNN, recurrent neural network.

| Tool (Year) | Description | Ref. |
|---|---|---|
| **Web servers for predicting pAs from DNA sequences** | | |
| Dragon PolyA Spotter (2012) | A web server for predicting 12 poly(A) motifs from human DNA sequences, using an artificial neural network and a random forest. | [60] |
| PolyApred (2009) | An SVM-based web server for predicting 13 poly(A) motifs in human, using sequence features of different types of nucleotide frequencies and binary pattern. | [58] |
| Polyadq (1999) | An early web server based on two quadratic discriminant functions for predicting AAUAAA/AUUAAA signals, using features encoded by position weight matrix. | [57] |
| **DL-based tools for predicting pAs from DNA sequences** | | |
| PASNet (2021) | A hybrid DL framework for identifying 16 poly(A) motifs in different species, which integrates gated convolutional highway networks with self-attention mechanisms. | [75] |
| SANPolyA (2020) | A self-attention deep learning model for predicting 18 poly(A) motifs in human and mouse. | [74] |
| HybPAS (2019) | A hybrid model for predicting 12 poly(A) motifs in human, using eight neural networks and four logistic regression models. | [73] |
| APARENT (2019) | The model was trained on isoform expression data from more than three million synthetic APA reporters. | [29] |
| DeepPASTA (2019) | A model based on CNN and RNN for predicting pAs from both sequence and RNA secondary structure. | [28] |
| DeeReCT-PolyA (2018) | A transferrable CNN model for recognition of 12 poly(A) motifs, which enables transfer learning across datasets and species. | [26] |
| DeepGSR (2018) | An approach based on CNN and one-hot features to predict genome-wide and cross-organism genomic signals and regions. | [27] |
| DeepPolyA (2018) | A model for predicting pAs in Arabidopsis with one-hot encoding features. | [71] |
| **Traditional ML-based tools for predicting pAs from DNA sequences** | | |
| PASS (2007) | A GHMM-based model for predicting pAs in plants. | [68] |
| Polya_svm (2006) | An SVM-based tool for predicting pAs using position-specific scoring matrices to score 15 *cis* regulatory elements. | [24] |
| **Methods for RNA-seq that rely on prior annotations of pAs** | | |
| QAPA (2018) | It compiles an expanded compendium of known pA annotations for identifying and quantifying pAs, which was suggested by Shah et al. [50] to be used in combination with pAs derived from 3′ seq or Iso-Seq. | [38] |

| | | |
|---|---|---|
| PAQR (2018) | It uses read coverage to segment 3′ UTRs at annotated pAs. | [86] |
| **Methods for RNA-seq that based on detecting changes in RNA-seq read density** | | |
| moutainClimber (2019) | It runs on a single RNA-seq sample and can recognize multiple transcription start sites or pAs. | [93] |
| APAtrap (2018) | It can detect all pAs along the 3′ UTR and can be used to improve 3′ end annotations. | [39] |
| TAPAS (2018) | It adopts a method originally used for time series data to detect change points, which was suggested have overall high performance in several benchmark studies [49, 50]. | [40] |
| DaPars (2014) DaPars2 (2018) | DaPars is probably the first and the most widely used tool for bulk RNA-seq and DaPars2 is its updated version. | [17, 90, 91] |
| **Methods for RNA-seq that based on ML models** | | |
| Aptardi (2021) | A multi-omics deep learning-based approach for predicting pAs by leveraging DNA sequences, RNA-seq, and the predilection of transcriptome assemblers. But its sensitivity may be low according to our preliminary test (Figure 3). | [98] |
| Terminitor (2020) | A DL-based model for three-label classification problem, which ddetermines a poly(A) cleavage site, a non-polyadenylated cleavage site, or non-cleavage site. | [97] |
| TECtool (2018) | It is based on transcriptome assembly and prior pA annotations, and can predict novel terminal exons. | [95] |
| **Methods for predicting pAs from scRNA-seq** | | |
| scDaPars (2021) | It is applicable to both full-length and 3′ tag scRNA-seq, which uses DaPars to infer pAs and may be slow for large scale scRNA-seq. | [45] |
| MAPPER (2021) | An annotation-assisted method for both bulk RNA-seq and 3′ tag scRNA-seq data, which incorporates prior pAs in the PolyA_DB for identifying pAs in 3′ UTRs and introns. | [105] |
| SCAPTURE (2021) | An annotation-assisted pipeline that implements a DL model to evaluate called peaks from 3′ tag scRNA-seq, using prior pAs from four databases for model training. | [106] |
| scUTRquant (2021) | An annotation-assisted method that incorporates a cleavage site atlas established from a mouse full-length Microwell-seq dataset of 400,000 single cells [108] for filtering pAs predicted from 3′ tag scRNA-seq. | [107] |
| SCAPE (2022) | A peak calling based method based on a probabilistic mixture model for identification and quantification of pAs in 3′ tag scRNA-seq by utilizing insert size information | [103] |
| ReadZS (2021) | A statistical approach to characterize read distributions that bypasses parametric peak calling and identify pAs from 3′ tag scRNA-seq. | [104] |
| scAPAtrap (2020) | A peak calling based method that incorporates poly(A) reads for genome-wide pA prediction from 3′ tag scRNA-seq. | [44] |
| Sierra (2020) | A splice-aware peak calling based method that can identify pAs in 3′ UTRs and introns from 3′ tag scRNA-seq. | [43] |
